# Population variability in social brain morphology: links to socioeconomic status and health disparity

**DOI:** 10.1101/2020.09.24.312017

**Authors:** Nathania Suryoputri, Hannah Kiesow, Danilo Bzdok

## Abstract

Health disparity across layers of society involves reasons beyond the healthcare system. Socioeconomic status (SES) shapes people’s daily interaction with their social environment, and is known to impact various health outcomes. Using generative probabilistic modeling, we investigated health satisfaction and complementary indicators of socioeconomic lifestyle in the human social brain. In a population cohort of ~10,000 UK Biobank participants, our first analysis probed the relationship between health status and subjective social standing (i.e., financial satisfaction). We identified volume effects in participants unhappy with their health in regions of the higher associative cortex, especially the dorsomedial prefrontal cortex (dmPFC) and bilateral temporo-parietal junction (TPJ). Specifically, participants in poor subjective health showed deviations in dmPFC and TPJ volume as a function of financial satisfaction. The second analysis on health status and objective social standing (i.e., household income) revealed volume deviations in regions of the limbic system for individuals feeling unhealthy. In particular, low-SES participants dissatisfied with their health showed deviations in volume distributions in the amygdala and hippocampus bilaterally. Thus, our population-level evidence speaks to the possibility that health status and socioeconomic position have characteristic imprints in social brain differentiation.

## Introduction

Most recently, the COVID-19 pandemic brought inequalities in socioeconomic status (SES) into sharp focus. Disproportionately worse health outcomes were reported in low-SES individuals (Patel et al. 2020). SES is a multifactorial construct that tracks dimensions of lifestyle, habitus, and world view, such as economic resources, occupational prestige and education attainment (Farah, 2017; Muscatell, 2018). A variety of studies showed that the “better off” an individual is as captured by common SES measures, the lower their risk to incur detrimental mental and physical health outcomes (Nancy E. Adler & Newman, 2002; Braveman et al., 2005).

In general, health disparities may be related to limited access to community resources or select lifestyle choices. Compared to high-SES individuals, low-SES individuals tend to have less adequate living conditions, higher incidences of smoking, alcohol consumption and obesity, less physical lifestyles, or have limited access to sufficient nutrition or health care services (N. E. Adler et al., 1994; Gwatkin, 2017; Sapolsky, 2004). Indeed, one study demonstrated systematic differences in social distancing behavior during the COVID-19 pandemic based on socioeconomic level (Weill et al. 2020). Using mobile device data, the results revealed high-income individuals to practice safe social distancing to a greater degree than low-income individuals. Such previous studies underscore how differences in one’s socioeconomic standing impact long-term health and life outcomes. Regular interaction with other people is also intimately related to our overall wellbeing. The experience of social isolation is linked to deteriorated health, and even shortened life expectancy (Bzdok & Dunbar, 2020; Cacioppo & Hawkley, 2003). One meta-analytic survey in ~300,000 individuals reported a shortage of social relationships to contribute to precipitated mortality, as well as risk factors including obesity, smoking and lack of exercise (Holt-Lunstad et al., 2010).

One of the earliest investigations on the interplay between health and SES comes from a longitudinal social epidemiological study with male British civil servants (Marmot et al., 1978). Mortality rates from coronary heart disease were negatively associated with a person’s position in the occupational hierarchy. Roughly three decades later, further social epidemiological studies reported increased risks of cancer, respiratory and gastrointestinal diseases and injuries among low-SES individuals (Mackenbach et al., 2003). In addition to higher mortality rates, low socioeconomic standing amplifies the risk for disability and psychiatric disorders such as major depressive disorder (Lorant et al., 2003). Low SES has even been linked to lower general intelligence (Yang et al., 2016), which in turn is related to poorer decision making of lifestyle choices and thus long-term health outcomes.

In contrast to objective SES measures such as yearly income, subjective or perceived indicators of SES reflect a person’s perspective on where they stand in the social hierarchy in comparison to others (Muscatell, 2018). The psychosocial nature of subjective SES indicators may also pinpoint an individual’s perceived well-being and personal experiences of deprivation better than objective SES measures (Brito & Noble, 2014; Singh-Manoux et al., 2007). Subjective SES measures have previously been found to be as valuable as objective SES indicators in predicting health outcomes (Singh-Manoux et al. 2005). For instance, a longitudinal study compared objective and subjective SES in forecasting health status. Objective SES was measured as one’s position in the occupational hierarchy and subjective SES was indexed as one’s perceived anchoring on the social ladder of society. The authors found in ~5500 middle-aged adults that over time, subjective SES predicted health status and health decline better than objective SES (Singh-Manoux et al. 2005). Moreover, a population health survey asked ~21,000 participants to identify their own anchoring in social network layers by indicating how satisfied they were with their finances (Li et al., 2018). The study showed people who self-identified as being in a lower social class were nine times more likely to report poor self-rated health than excellent health, compared to higher-SES individuals (Li et al., 2018). Thus, objective and subjective measures of SES may together capture complementary aspects and provide a clearer understanding of an individual’s embedding in the societal hierarchy.

Varying levels of SES were shown in neuroimaging studies to reveal different manifestations in brain structures important for health. These differences in neural architecture have been reported to be detectable as early as the age of ~8 years (Farah, 2017; Hackman & Farah, 2009). In adults, a structural brain-imaging study observed smaller amygdala and hippocampal volumes for participants experiencing financial hardship compared to participants more financially stable (Butterworth et al. 2012). These two limbic regions are involved in the physiological regulation of stress, and are also known to be neural substrates with important implications for social interaction processes (e.g., Bickart et al., 2011).

The stress of experiencing financial hardship may have downstream consequences on influencing limbic grey matter volume via functional alterations in cortisol release through the hypothalamic-pituitary-adrenocortical axis (Gianaros et al. 2007; Butterworth et al. 2012). Indeed, a previous neuroimaging study investigated the link between socioeconomic disadvantage and cortical brain structure in n=~450 middle-aged adults (Gianaros et al. 2017). The authors found socioeconomic disparity, indexed by a composite measure of households under the poverty level and receiving public assistance, education level, employment status and household income, to be closely linked to reduced grey matter volume in regions of the frontal and temporal lobes (Gianaros et al., 2017). Notably, some of the brain regions reported by Gianaros and colleagues are also known to be implicated in several fundamental social processes such as the dorsomedial prefrontal cortex, fusiform gyrus, and temporo-parietal junction (Alcalá-López et al. 2018; Mitchell 2009; Pelphrey et al. 2004). Gianaros and colleagues suggest socioeconomic adversity may have widespread manifestations in neural architecture through one’s health status.

Exposure to stressful or negative experiences may explain the observed SES-related discrepancies in features of brain architecture. The experience of chronic stress is known to increase the neuroendocrine response and thus amplify the risk for stress-related diseases (N. E. Adler et al., 1994; Sapolsky, 2004, 2005). In line with this contention, a previous brain imaging study found low-SES individuals to show enhanced activity in the dmPFC in response to social threat, compared to participants higher up on the social hierarchy (Muscatell et al., 2016). Furthermore, neural activity changes in the dmPFC were found to mediate the relationship between social status and increases in inflammation, a marker for illness (Muscatell et al., 2016). Thus, these previous studies provide evidence that differences in one’s socioeconomic circumstances may ultimately relate to longevity, and hint to neural representations that may underpin SES-health gradients.

Despite the intimate relationship between SES and health, accumulating evidence shows that more social ties and tighter connections may act as a buffer against potential illness. For example, a population-sized longitudinal study (n=~5000) found that belonging to more social groups protected against future depression in healthy participants (Cruwys et al., 2013). Notably, the authors also found that being more socially engaged alleviated and prevented relapse of existing depression symptoms in depressed participants (Cruwys et al., 2013). Similarly, a behavioral study administered flu vaccines to participants in an effort to assess the relationship between illness and the strength of social ties (Pressman et al. 2005). The authors found lonely individuals with few friends to show the most reduced immune response compared to individuals with larger social networks, and those better surrounded (Pressman et al., 2005). Together, these studies converge on the contention that cultivating interpersonal relationships may provide protective advantages for long-term health.

In summary, earlier findings hint at how different levels of socioeconomic status may interfere with long-term health. Previous neurobiological evidence has also revealed SES-related manifestations in several brain regions important for health. To complement these aforementioned investigations, our study probed the extent to which health satisfaction and two measures of SES are reflected in social brain morphology. In contrast to previous neuroimaging studies with small sample sizes, we used the uniformly acquired 10,000 participant release from the UK Biobank initiative - the world’s largest available biomedical database. For anatomical guidance into the human social brain, we employed a recently available social brain atlas. These topographical definitions included four brain networks at increasing hierarchical level and their 36 brain regions that were consistently implicated in social and affective processes (Alcalá-López et al., 2018). To zoom in on variation in social brain morphology, our analyses implemented a probabilistic modelling perspective. We purpose-designed a framework that explicitly modelled the extent of similarity and divergence between health-specific social brain volume patterns in the context of objective (i.e., income) and subjective (i.e., financial satisfaction) markers of SES. In two complementary analyses, we thus modelled the 36 social brain regions with their relation to four participant subgroups: simultaneously examining participants unsatisfied versus satisfied with their health in a low-versus high-SES indicator.

## Methods

### Data resources

The UK Biobank is a prospective epidemiology resource that offers extensive behavioral and demographic assessments, medical and cognitive measures, as well as biological samples in a cohort of 500,000 participants recruited from across Great Britain (Sudlow et al., 2015). This openly accessible population dataset aims to provide multimodal brain-imaging for 100,000 individuals to be completed in 2022 (Miller et al., 2016). The present study was based on the data release providing brain-imaging from 10,129 individuals to detail population variation in greymatter morphology of the social brain as measured by T1-weighted structural magnetic resonance imaging (MRI). All participants were scanned at the same assessment center. Improving comparability and reproducibility, our study profited from uniform data preprocessing pipelines that allow for easier comparability to other and future UKBB population studies (Alfaro-Almagro et al., 2018). The population sample included 47.6% males and 52.4% females, aged 40-69 years when recruited (mean age of ~55 years, standard deviation (SD) ± 7.5 years). The present analyses were conducted under UK Biobank application number 23827. All participants provided informed consent to participate (http://biobank.ctsu.ox.ac.uk/crystal/field.cgi?id=200).

For the present population neuroscience study, three health and lifestyle-related summary measures from the UKBB were considered: i) health satisfaction, ii) income as an objective measure of SES and iii) financial satisfaction as a subjective measure of SES. For the SES measure of income (UKBB ID: 738), participants were divided into the high-income subgroup if their yearly income exceeded £100,000, while the remaining participants were included in the low-income group. For the SES measure of financial satisfaction (UKBB ID: 4581), participants were asked “In general how satisfied are you with your financial situation?”. Participants who rated themselves as extremely or very happy with their financial situation were included in the financially satisfied subgroup, while the remaining participants were included in the financially unsatisfied subgroup. Lastly, for the health satisfaction measure (UKBB ID: 4548), participants were asked “In general how satisfied are you with your health?”. Participants who rated themselves as extremely or very happy with their health were included in the subgroup of people feeling healthy, while the remaining participants were included in the subgroup of people feeling unhealthy.

### Brain-imaging preprocessing procedures

Identical MRI scanners (3T Siemens Skyra) were used at the same imaging site with the same acquisition protocols and standard Siemens 32-channel radiofrequency receiver head coils. To protect the anonymity of the study participants, brain-imaging data were defaced and any sensitive information from the header was removed. Automated processing and quality control pipelines were deployed (Miller et al. 2016). To improve homogeneity of the brain-imaging data, noise was removed by means of 190 sensitivity features. This approach allowed reliable identification and exclusion of problematic brain scans, such as scans with excessive head motion.

The structural MRI data were acquired as high-resolution T1-weighted images of brain anatomy using a 3D MPRAGE sequence at 1 mm isotropic resolution. Preprocessing included gradient distortion correction, field of view reduction using the Brain Extraction Tool (Smith, 2002) and FLIRT (Mark Jenkinson et al., 2002; M. Jenkinson & Smith, 2001), as well as nonlinear registration to MNI152 standard space at 1 mm resolution using FNIRT (Andersson et al., 2007). To avoid unnecessary interpolation, all image transformations were estimated, combined and applied by a single interpolation step. Tissue-type segmentation into cerebrospinal fluid, grey matter (GM) and white matter was applied using FAST (FMRIB’s Automated Segmentation Tool, Zhang et al. 2001) to generate full bias-field-corrected images. Analyses in the present study capitalized on the ensuing GM maps. In turn, SIENAX (Smith et al., 2002) was used to derive volume measures normalized for head sizes. The ensuing adjusted volume measurements represented the amount of grey matter corrected for individual brain sizes.

### Social brain atlas definition

Our study benefited from a recent quintessential definition of the social brain in humans. This topographical atlas was derived by a large-scale quantitative synthesis of a diversity of functional MRI (fMRI) findings from thousands of experimental studies involving 22,712 individuals (Alcalá-López et al., 2018). 36 volumes of interest were previously identified with consistent neural activity increases during a wide assortment of social and affective tasks. The 36 data-derived target locations were also shown to be connectionally and functionally segregated into i) a visual-sensory network, ii) a limbic network, iii) an intermediate network, and iv) a higher-associative network. This existing knowledge of how the social brain regions relate to major brain networks at different hierarchical processing levels was incorporated in our modeling approach (cf. below).

The topographical specificity of our targeted analyses was thus enhanced by guiding brain volume extraction by the 36 volumes of interest, each associated with one of four social brain networks. Neurobiologically interpretable measures of grey-matter volume were extracted in the ~10,000 participants by summarizing whole-brain anatomical maps guided by the topographical compartments of the social brain (Kernbach et al., 2018; Miller et al., 2016). We applied a smoothing filter of 5mm FWHM to the participants’ structural brain maps to homogenize local neuroanatomical differences (Kiesow et al. 2020). Local quantities of social brain morphology were then extracted as 36 average volume measures per participant. Grey-matter measures were averaged in spheres of 5mm diameter around the consensus location from the social brain atlas, averaging the preprocessed, tissue-segmented, and brain-size-adjusted MRI signal (cf. above) across the voxels belonging to a given target region (cf. Supl. Table 1 for stereotaxic MNI coordinates for each of the social brain atlas regions). The grey-matter information from all voxels belonging to a particular atlas region were added up and divided by the total number of region voxels. These operations returned a single representative measure for the mean grey-matter volume in the particular brain region. Note that using a smaller sphere diameter of 2.5mm or bigger spheres of 7.5mm yielded virtually identical results. This way of engineering morphological region summaries yielded 36 volume brain variables per participant, as many as the total number of social brain regions, which were subsequently z-scored by centering to zero mean and unit-variance scaled to one. These commonly employed estimates of population brain volume variability (Kernbach et al., 2018; Miller et al., 2016) in social brain anatomy served as the basis for all subsequent analysis steps.

All regions of interest used in this study are available online for transparency and reuse at the data-sharing platform NeuroVault (http://neurovault.org/collections/2462/).

### Probabilistic hierarchical modeling of volume variation in the social brain

To jointly model the population distribution of brain volume effects underlying the relationship between subjective health and socioeconomic contexts, we designed and carried out a Bayesian hierarchical regression analysis (Gelman et al., 2013), building on our previous research (Bzdok et al. 2020; Kiesow et al. 2020; Bzdok et al. 2017). Traditionally, classical null-hypothesis-testing approaches provide p-values against the null hypothesis of no effect in the data. Instead, our goal was to equally consider the outcomes of the presence vs. absence of health satisfaction and socioeconomic distinctions in human social brain regions in all parts of our modeling approach. We thus “let the data speak for themselves” by directly interrogating the population uncertainty intervals of volume parameter distributions in their relation to SES and health, rather than restricting attention to differences in mean volume alone.

The probability model with parameters that vary by network followed the following form:

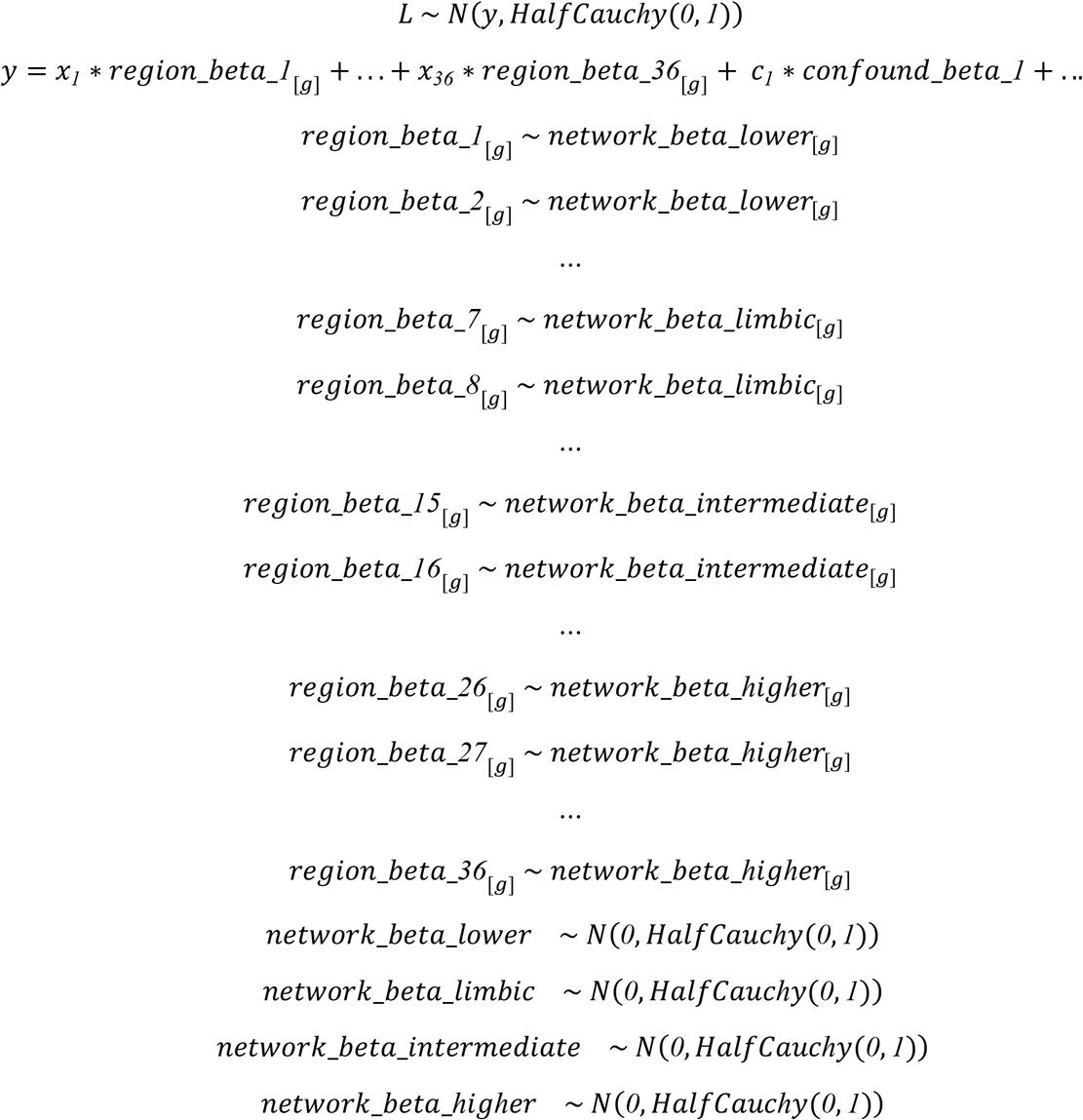

where *x_i_* denote the brain volumes for all 36 brain regions of the social brain atlas (z-scored across participants), *y* denotes the age of the participants, Gaussian-distributed hyper-beta coefficients underlying network-level volume variation jointly informs beta coefficients at the region level for each participant subgroup *g*. Variance that could be explained by the nuisance variables *c_i_* of no interest body mass and head size was accounted for as potential confounds (Kernbach et al., 2018; Miller et al., 2016). Additionally incorporating height and weight as covariates of no interest in the analyses did not change the results and led to the same conclusions. For each analysis, each participant was assigned to one of four subgroups *g* that indicated stratification of our population sample into two levels of health satisfaction, namely feeling healthy and feeling unhealthy, along with the presence of two socioeconomic measures, those with a high or low income and perceived financial satisfaction or dissatisfaction. In this way, we could get the most out of our rich sample by borrowing statistical strength between clusters of individuals in our population cohort through interlocking of their model coefficients. Parameters of the within-subgroup regressions were placed at the bottom of the model architecture were modeled themselves by the hyper-parameters of the across-subgroup regression to pool information across batches of variance components. We could thus provide more detailed quantitative answers to questions about morphological differences of the social brain by a joint model estimation profiting from several sources of population variation.

Approximate posterior inference was achieved by Markov Chain Monte Carlo (MCMC) using PyMC3 in Python (https://github.com/pymc-devs/pymc3), which sampled in a random walk towards the target posterior distribution. In 5,000 draws, the approximate parameter distributions were improved at each step in the sense of converging to the target distribution. At each step of the MCMC chain, the entire set of parameter values were estimated to be jointly credible given the data. A range of explanations for the relation between health status and SES level were browsed through by obtaining multiple plausible settings of model parameters that could have led to the observed data. We searched through configurations of parameters as an efficient way of exploring the important part of the parameter space, and then averaged the resulting models according to their quality. In particular, we dropped the 4,000 first samples from the chain because 1) the chain had probably not yet reached stationarity and 2) this step reduced dependence on the starting parameter values. After tuning the sampler for 4,000 steps, we drew 1,000 samples from the joint posterior distribution over the full set of parameters in the model for analysis. Proper convergence was assessed by ensuring Rhat measures (Gelman et al. 2013) stayed below 1.02.

The present modeling strategy has at least three advantages (Gelman et al., 2013). First, we could directly quantify how the tail area uncertainty the region volumes of social brain vary as a function of socioeconomic traits and health outcome. Additionally, the usual problem of multiple comparisons is automatically addressed because hierarchical modeling can find large differences as a byproduct of searching through many parameter constellations (Gelman et al., 2013; Kruschke, 2011). Second, our probabilistic hierarchical regression was aware of the meaningful stratification in our dataset by simultaneously estimating within-subgroup variation and between-subgroup variation from the behavioral and brain data simultaneously. Third, appreciating the existing hierarchical structure in our data is known to stabilize posterior estimation and guard against overfitting to noise.

### Scientific computing implementation

Python was selected as the scientific computing engine. Capitalizing on its open-source ecosystem helps enhance replicability, reusability, and provenance tracking. The scikit-learn package (Pedregosa et al., 2011) provided efficient, unit-tested implementations of state-of-the-art machine learning algorithms (http://scikit-learn.org). This general-purpose machine-learning library was interfaced with the nilearn library (Abraham et al., 2014) for design and efficient execution of neuroimaging data analysis workflows (http://github.com/nilearn/nilearn). 3D visualization of brain maps was performed using PySurfer (https://pysurfer.github.io/), and data plots were generated by Seaborn (https://seaborn.pydata.org/). Probabilistic hierarchical modeling and MCMC sampling was implemented as symbolic computation graphs in the PyMC3 framework (https://github.com/pymc-devs/pymc3). All analysis scripts that reproduce the results of the present study are readily accessible to and open for reuse by the reader (http://github.com/banilo/to_be_added_later).

## Results

The present population neuroscience study explored the intricate relationship between health disparity and socioeconomic environments in the context of social brain morphology. To probe the relationship between SES and health indicators, we estimated two probabilistic hierarchical models. One analysis focused on health satisfaction and subjective SES (i.e., financial satisfaction), while the other one focused on health satisfaction and objective SES (i.e., income). We designed these fully probabilistic, generative models to simultaneously estimate divergences in regional grey matter volume between subgroups at the hierarchical network level and at the region level. Thus, each quantitative analysis investigated health satisfaction and a different measure of SES in ~10,000 UK Biobank participants to directly estimate posterior probability distributions of social brain volume at the population-level.

For each analysis, the participants were divided into four subgroups as combinations of SES and health levels. In the first analysis, subgroups were as follows: i) low health satisfaction and low financial satisfaction, ii) low health satisfaction and high financial satisfaction, iii) high health satisfaction and low financial satisfaction, and iv) high health satisfaction and high financial satisfaction. Analogously, in the second analysis, subgroups were as follows: i) low health satisfaction and low income, ii) low health satisfaction and high income, iii) high health satisfaction and low income, and iv) high health satisfaction and high income. Henceforth, the term “volume effect” refers to parameter distributions of the probabilistic model that reflect effects in grey matter volume variation in a population neuroscience context (cf. Materials and Methods). In this way, we were able to ask “How certain are we that the grey matter volume of an individual social brain region and its brain network are similar or divergent among subpopulations of different health status and socioeconomic standing?”.

Overall, at the network-level of the modeling results, participants with a higher quality of health showed similar volume divergences in regions of the visual-sensory network and higher associative network. However, regions of the limbic network showed diverging results in our analyses involving the two SES measures. For instance, in the context of income, we identified population volume effects in the limbic network predominantly in the subgroups of participants unhappy with their health condition. Conversely, we found individuals satisfied with their health to show volume parameter dispersions in the limbic network as a function of financial satisfaction. Intermediate-network-wide volume effects were detected with respect to household income mostly for the subset of individuals with a better perceived quality of health. However, volume effects were observed in the intermediate network for individuals with a lower perceived health standing, in relation to financial satisfaction.

### Health satisfaction and financial satisfaction in social brain structure

At the highest level of hierarchical networks in the social brain atlas, we observed strong volume effects among the participant subgroups with a lower perceived quality of health. Notably, individuals expressing dissatisfaction with their financial situation showed volume effects in the dmPFC, TPJ_R and MTG_L that deviated from the population volume distributions of participants more satisfied with their finances (Fig. 1; see Supl. Table 1 for abbreviations). In contrast, among the subgroups of participants happy with their health, individuals also content with their finances showed a divergent trend in volume variation in the PCC, Prec and pMCC compared to those with a lower perceived SES (see Supl. Table 2 for full results regarding the mean of the posterior distribution and highest posterior density interval (HPDI) covering 95% uncertainty). However, interindividual variation of grey matter volume in relation to health- and financial satisfaction was reflected in participants in different ways in the TPJ_L and bilateral TP (Fig. 1). Specifically, in the TPJ_L, a pronounced volume effect was observed for individuals with poor perceived health quality and low financial satisfaction, compared to participants with the same health status but feeling more satisfied with their finances. We observed in the TPJ_L an additional volume effect among the subgroups of participants with good health status. Specifically, those satisfied with their finances deviated in volume from participants with a lower level of financial satisfaction. Conversely, in the bilateral TP, in higher health satisfaction, individuals happy with their finances showed divergent volume distributions compared to participants less satisfied with their finances. Likewise, we detected evidence for anatomical divergence in the bilateral TP among participants with poor health in the context of perceived SES. In summary, pronounced grey matter volume effects in regions of the higher associative network were observed among both the healthy and unhealthy subgroups, where deviations in volume were most apparent for different levels of financial satisfaction.

**Figure 1.**
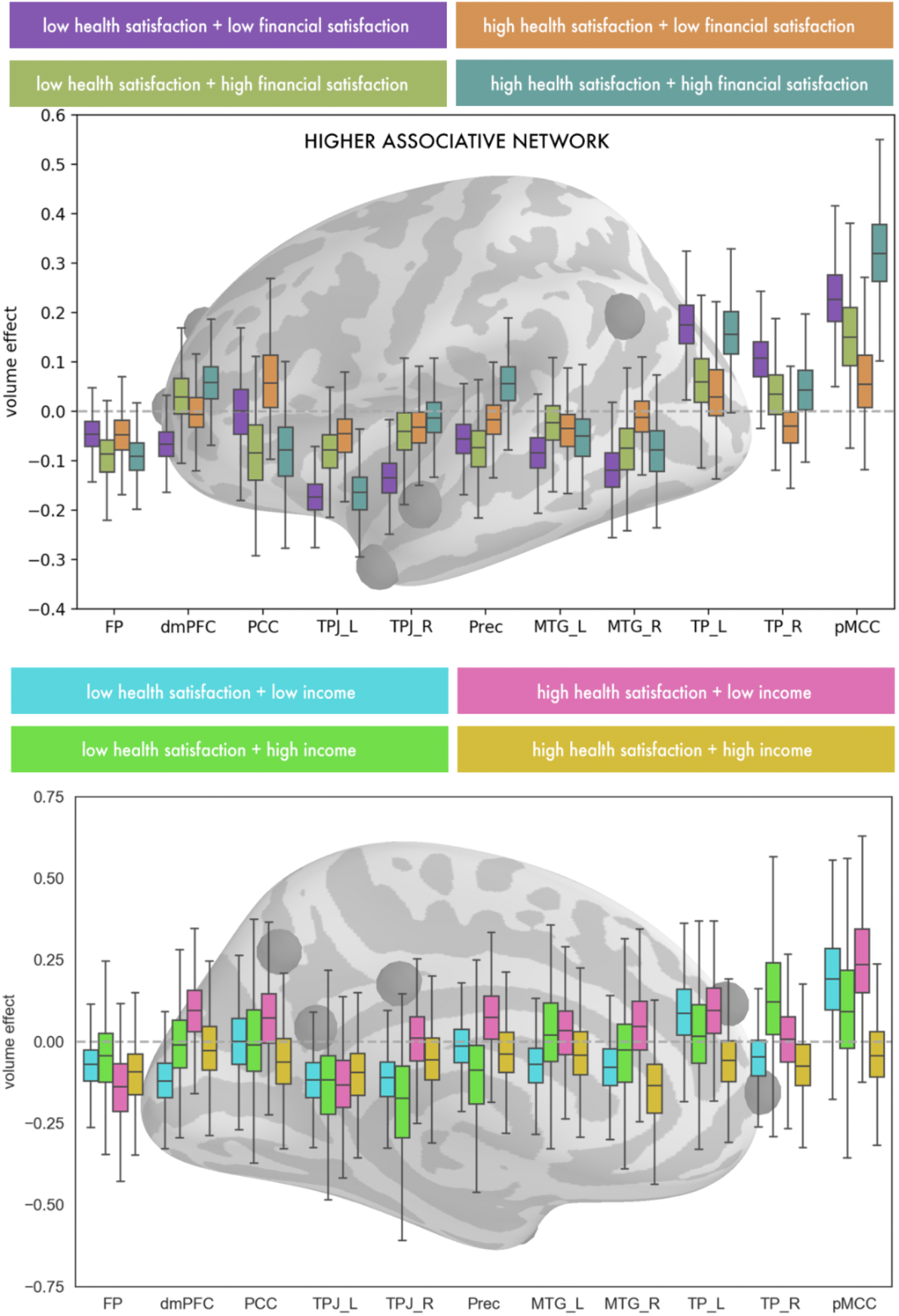
Good health status and high subjective socioeconomic standing are linked to higher associative regions related to perspective taking. Marginal posterior population distributions as depicted in the boxplots reveal the extent to which SES measures and health satisfaction vary for a particular subgroup for each higher associative region (y-axis). The top panel depicts volume effects as a function of health satisfaction and financial satisfaction. The bottom panel shows the results for health satisfaction and household income. These population-level findings suggest a degree of health-specific sensitivity among most higher associative social brain regions as a function of socioeconomic standing. For abbreviations see Supplementary Table 1.

In the intermediate network, volume effects related to financial satisfaction were observed especially among participants feeling unsatisfied with their health. Specifically, in the AI_R and SMA_R, strong volume effects were observed for the subset of UKBB participants unhappy with their finances compared to those more satisfied with their financial situation (Fig. 2). This pattern in population volume variation was associated with largely overlapping parameter distributions in the AI_R and SMA_R among the subgroups of participants feeling more fulfilled with their health. However among the subgroups of participants with a high health rating, individuals with low financial satisfaction showed incongruent volume distributions in the SMG_R and Cereb_L compared to participants with a better financial situation (Fig. 2). This pattern in population volume variation in these intermediate network regions was not observed among the subgroup of participants with low health satisfaction. To summarize findings in the intermediate network, grey matter volume effects were most pronounced for individuals with a lower financial satisfaction, in both contexts of good and poor health.

**Figure 2.**
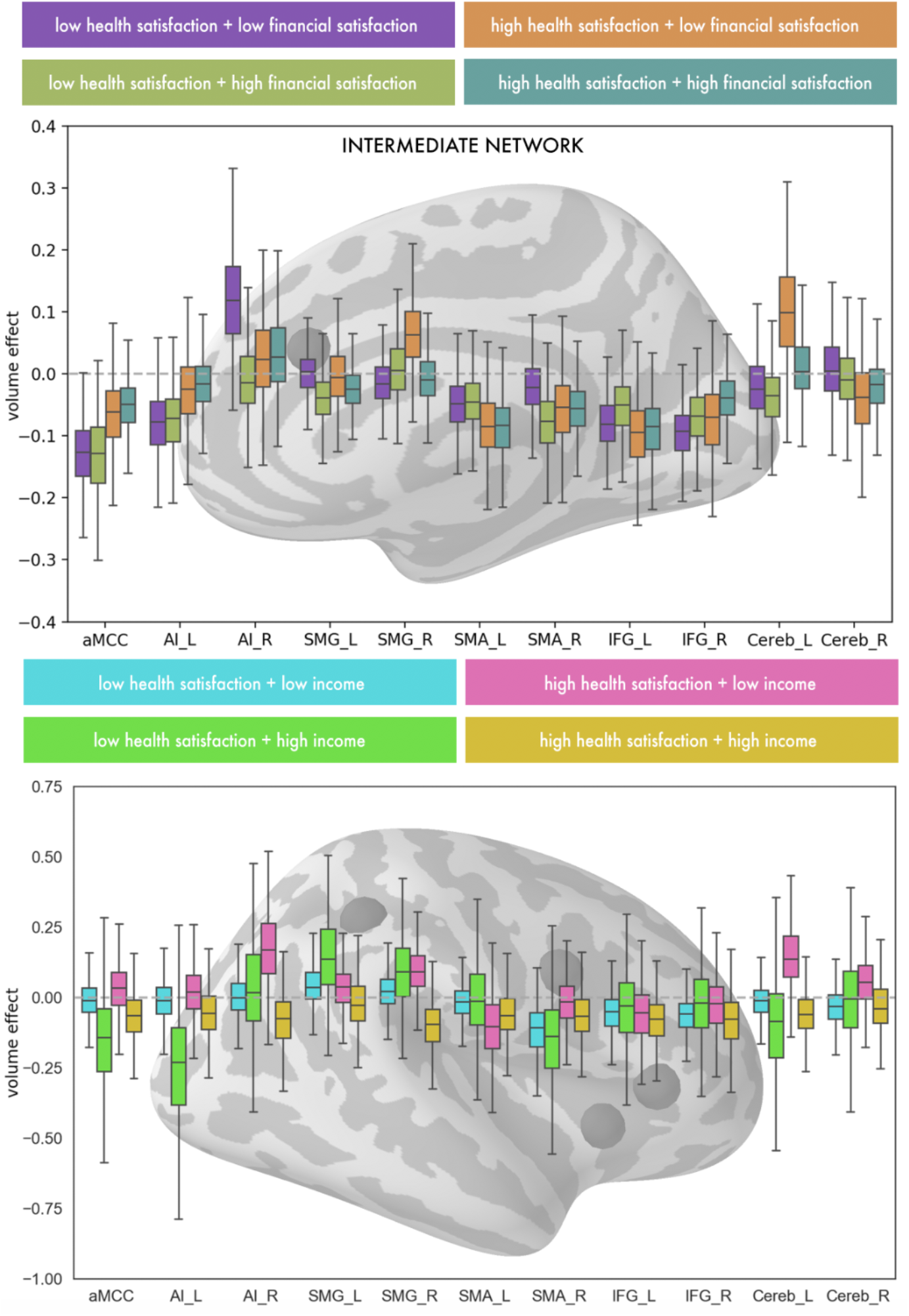
Good health status is linked to intermediate network regions in participants with low socioeconomic standing. Marginal posterior population distributions as depicted in the boxplots reveal the extent to which SES measures and health satisfaction vary for a particular subgroup for each higher associative region (y-axis). For each intermediate social brain region, a given set of volume effects represent a subgroup combination of low- or high-SES level and low or high health status. The top panel depicts the population results for health satisfaction and financial satisfaction. The bottom panel shows the results for health satisfaction and household income. These results highlight similar volume effects among participants expressing satisfaction with their health in both contexts of subjective financial satisfaction and reported yearly income. For abbreviations see Supplementary Table 1.

In regions of the limbic network, volume effects dependent on financial satisfaction became apparent among the subgroups of UKBB participants content with their health. Among the examined participants who expressed satisfaction with their health, those unhappy with their finances showed stronger volume effects especially in the AM_R and bilateral NAC compared to individuals satisfied with their financial situation (Fig. 3). In contrast to the observed volume effects linked to higher subjective physical well-being, we found volume effects in the limbic HC_L and vmPFC for those in poor subjective health (Fig. 3). Among these participants unsatisfied with their health, those with low financial satisfaction showed population parameter distributions that were incongruent from participants more satisfied with their level of SES. As such, the volume effects observed in the limbic network showed the most substantial deviations in the reward-related NAC and emotion-related AM.

**Figure 3.**
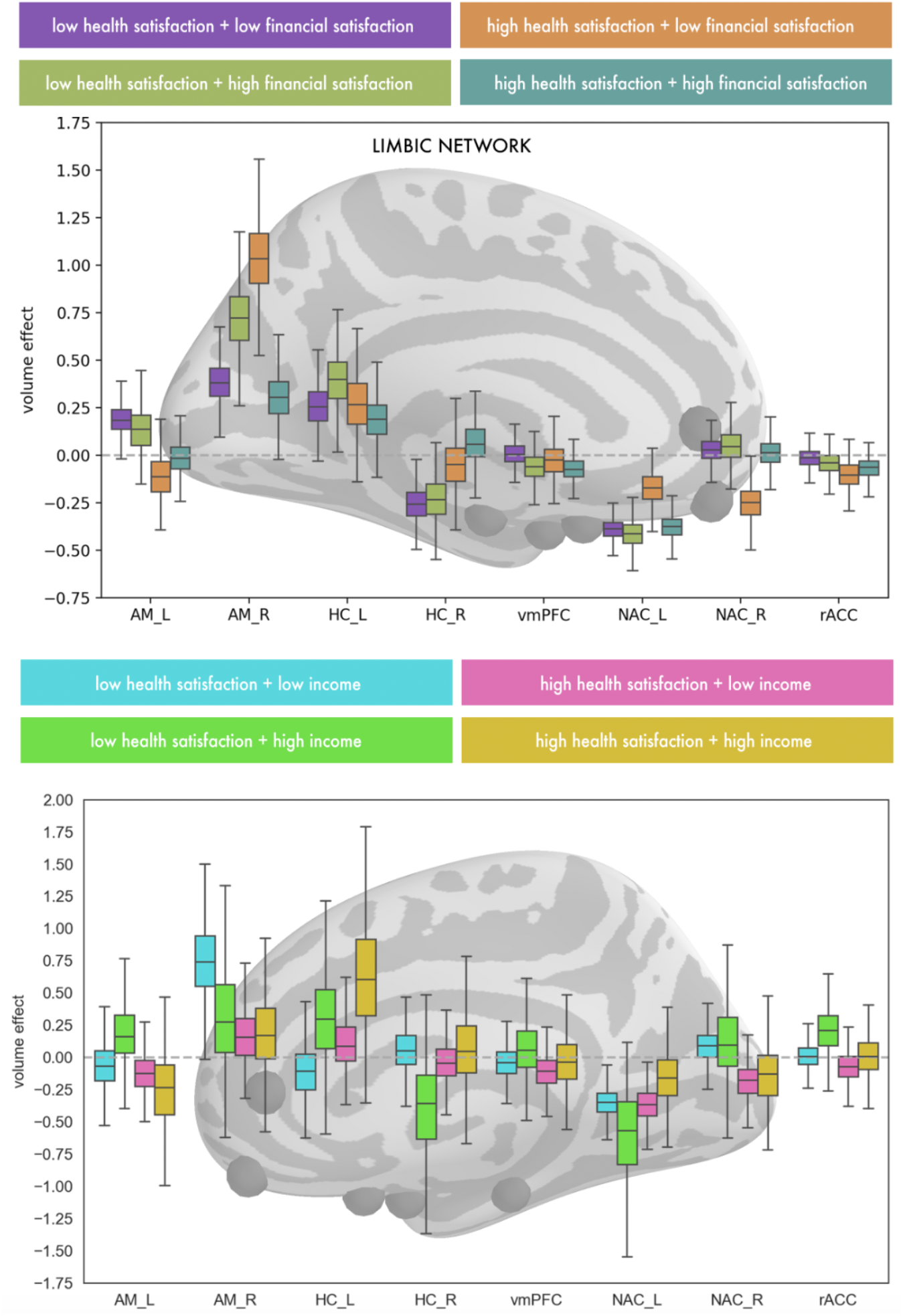
Low financial satisfaction is related to limbic regions for participants in good health. Marginal posterior population distributions as depicted in the boxplots reveal the extent to which SES measures and health satisfaction vary for a particular subgroup for each higher associative region (y-axis). For each limbic social brain region, a given set of volume effects represent a subgroup combination of low- or high-SES level and low or high health status. The top panel depicts the population results for health satisfaction and financial satisfaction. The bottom panel shows the results for health satisfaction and household income. These results reveal interindividual differentiation in limbic grey matter volume in the context of objective and subjective social standing and health disparity. For abbreviations see Supplementary Table 1.

Lastly, regarding social brain regions closest to sensory input processing, population parameter distributions of our model indicated volume effects in the majority of visual-sensory regions. These effects were especially apparent among the subgroups of participants expressing satisfaction with their health. Specifically, for participants in good perceived health, participants feeling unfulfilled with their finances predominantly showed volume deviations in the FG_R, bilateral pSTS, and MT/V5_L, compared to participants with high financial satisfaction (Fig. 4). In contrast, in participants with poor perceived health, different satisfaction levels with one’s finances were associated with more overlapping volume parameter distributions in these visual-sensory social brain regions. However, in the context of low health satisfaction, participants happier with their financial situation showed a pronounced volume effect in the MT/V5_R, compared to the examined participants unsatisfied with their SES level. In sum, interindividual variability in region volumes of the visual-sensory network was most pronounced for individuals with high health satisfaction, but unhappy with their financial situation.

**Figure 4.**
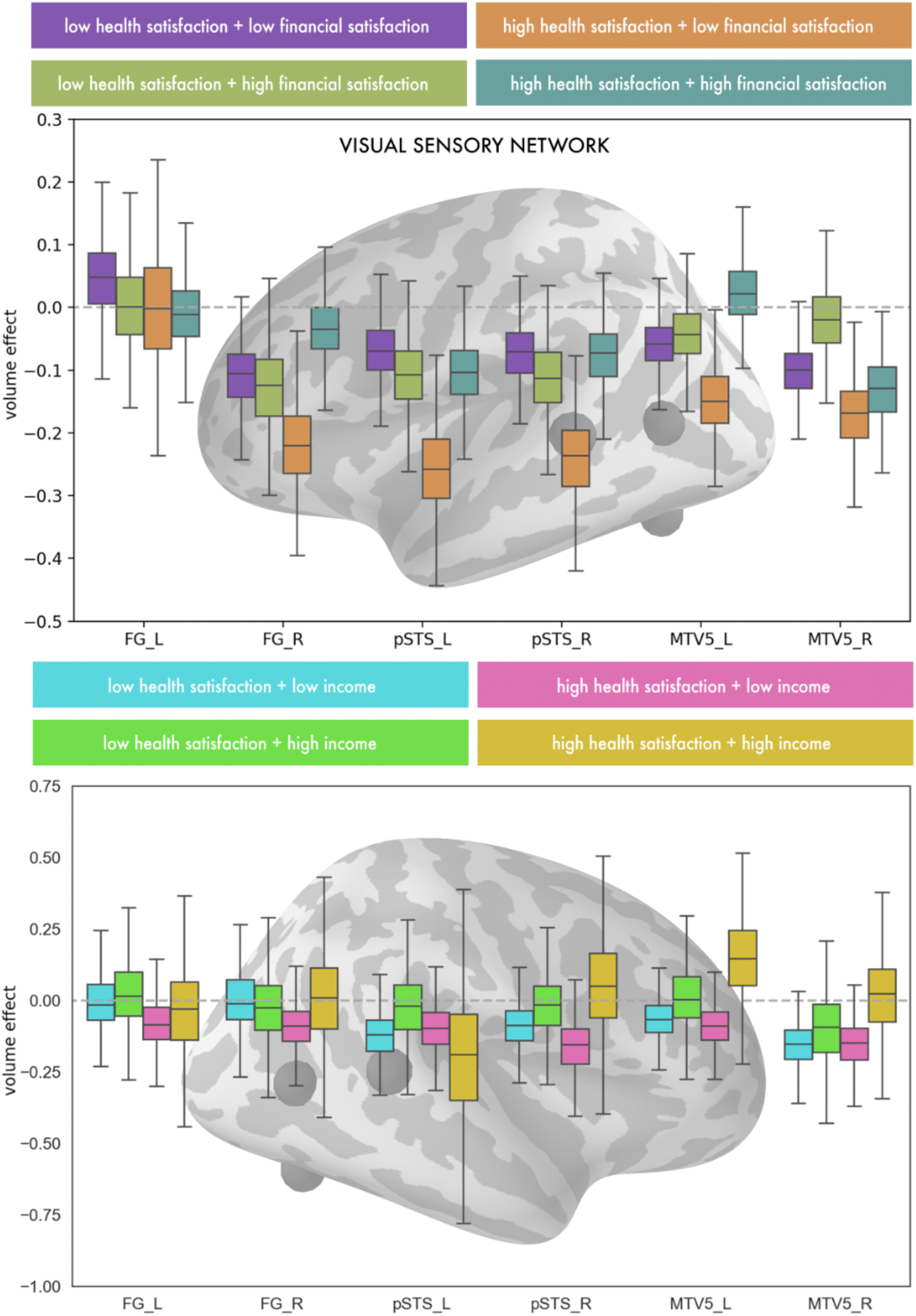
Good health status is linked to sensory network regions as a function of income and financial satisfaction. Marginal posterior population distributions as depicted in the boxplots reveal the extent to which SES measures and health satisfaction vary for a particular subgroup for each higher associative region (y-axis). For each intermediate social brain region, a given set of volume effects represent a subgroup combination of low- or high-SES level and low or high health status. The top panel depicts the population results for health satisfaction and financial satisfaction. The bottom panel shows the results for health satisfaction and household income. These results suggest health-specific sensitivity among most visual-sensory social brain regions as a function of differing objective and subjective socioeconomic environments. For abbreviations see Supplementary Table 1.

In summary, we show health satisfaction and a subjective measure of SES to show pronounced volume effects in all four hierarchical networks. Overall, regions of the higher associative network, limbic, and visual-sensory network showed particularly strong volume effects compared to regions of the intermediate network. Notably, divergences in social brain volume were typically observed for participants who were satisfied with their health.

### Health satisfaction and income in social brain structure

Regarding our analysis based on an objective measure of SES, in the higher-associative network of the social brain, pronounced volume effects became evident among participants with a higher perceived quality of health. Large population volume effects were observed in the Prec, MTG_R, TP_L, PCC and pMCC for individuals earning a high, as opposed to low, income (Fig. 1). However, for participants with low health satisfaction, different income levels were associated with more overlapping volume parameter distributions in these higher associative regions (see Supl. Table 3 for full results regarding the mean of the posterior distribution and highest posterior density interval covering 95% uncertainty). However, our model showed evidence for income-related structural divergences in the MTG_L and TP_R in the context of low health satisfaction (Fig. 1). In these two higher-associative regions, participants with poor subjective health and a low income deviated in grey matter volume from participants of the same perceived health status, but with a higher income. Similarly, we linked interindividual differences in both health and SES to pronounced volume effects in the dmPFC. Specifically, among UKBB participants with a lower health satisfaction, those also earning a low yearly wage showed population volume effects, compared to participants with a higher income. Similarly, for participants with good health, those earning a lower wage showed a divergent trend in dmPFC volume compared to participants with a higher wage. In summary, interindividual variation in grey matter volume was mostly evident in regions of the higher-associative network for the subset of healthy participants earning a higher income.

Parallelling the grey matter volume findings observed in the higher associative network, we also observed salient volume effects in the intermediate network among the subgroups of UKBB participants with a high health rating. Notably, incongruent volume dispersions were observed in healthy participants with high versus low income in the AI_R, SMG_R and Cereb_L (Fig. 2). This pattern of income-related volume effects was not observed among UKBB participants with low health satisfaction. However, in low-health participants, we observed a divergent trend in income-dependent volume distributions in the aMCC and AI_L as a function of yearly income (Fig. 2). As such, interindividual differences in income related to a variety of volume effects in the intermediate network.

A degree of income-related volume effects also became apparent in several regions of the limbic network, especially among participants expressing inadequate health in the bilateral AM, rACC and HC_R (Fig. 3). UKBB participants with a lower perceived quality of health and with a low income showed population parameter distributions in these limbic regions that were incongruent with higher income participants of the same poor health status. In contrast, in the context of high health satisfaction, different income levels were associated with overlapping posterior population distributions. However, among participants who felt in better shape, volume effects in the NAC_L were observed for high-income, as compared to low-income participants (Fig. 3). However, health- and income levels were expressed differently in grey matter volume variation in the HC_L. Specifically, low-health participants with low income showed the larger volume deviation in the HC_L compared to high-income participants. Conversely, participants with high health satisfaction, high-income participants showed volume deviations in the HC_L compared to low-income participants. Together, we observed morphological divergences in regions of the limbic system mostly in participants with a lower perceived health as a function of household income.

In the visual-sensory network in turn, several prominent volume effects related to income were observed in these brain regions of the lowest tier of the processing hierarchy. In particular, high-health participants revealed volume effects linked to income in the pSTS_R and bilateral MT/V5 (Fig. 4). In these regions, healthy individuals with a higher SES status showed population volume effects that diverged from the subset of UKBB participants with a lower SES status, but with the same perceived quality of health. For the groups of participants with a lower health status, overlapping volume distributions were mostly observed in these regions. Together, interindividual variation in regions of the visual-sensory network were mostly evident among participants with a higher health status.

Summarizing region effects related to our objective measure of SES, we observed the most frequent income-related divergences in population parameter distributions in the subgroups of individuals happier with their health. Specifically, within the subgroup of people with high health satisfaction, income-level related divergences in volume effects were shown to be similar in regions of the higher associative, intermediate and visual-sensory networks. Whereas in the limbic network, volume effects were most apparent among individuals unsatisfied with their health.

### Network-wide effects of health satisfaction and socioeconomic standing

To reiterate, our social brain atlas also contains four target brain networks that encompass the 36 target social brain regions (Fig. 5). In addition to modeling region-specific volume effects, our probabilistic framework simultaneously quantified brain network-wide volume effects as a function of health and SES. Income-related volume effects were observed in the two highest tiers of the social brain network hierarchy for individuals with high health status. In the higher associative network, participants earning a high yearly income deviated in population network volume compared to those of the same health status, but earning a lower yearly wage (Fig. 5). The volume effect in the higher associative network was paralleled by our results in the intermediate network. Individuals happy with their health and earning a high income showed volume deviations, compared with participants of the same high health status but earning a lower income. Among the subgroups with a low health status, participants with low and high incomes were more congruent in posterior parameter distributions that captured volume effects in the entire intermediate network. In contrast, in the context of subjective SES, we observed a network volume effect in the lowest tier of the processing hierarchy among the subgroups of participants expressing contentment with their health. Participants discontent with their finances showed incongruent visual-sensory network volume effects relative to participants with a better financial situation. Such coherent patterns of network-wide divergence were not observed in the other networks of our social brain atlas in the context of financial satisfaction.

**Figure 5.**
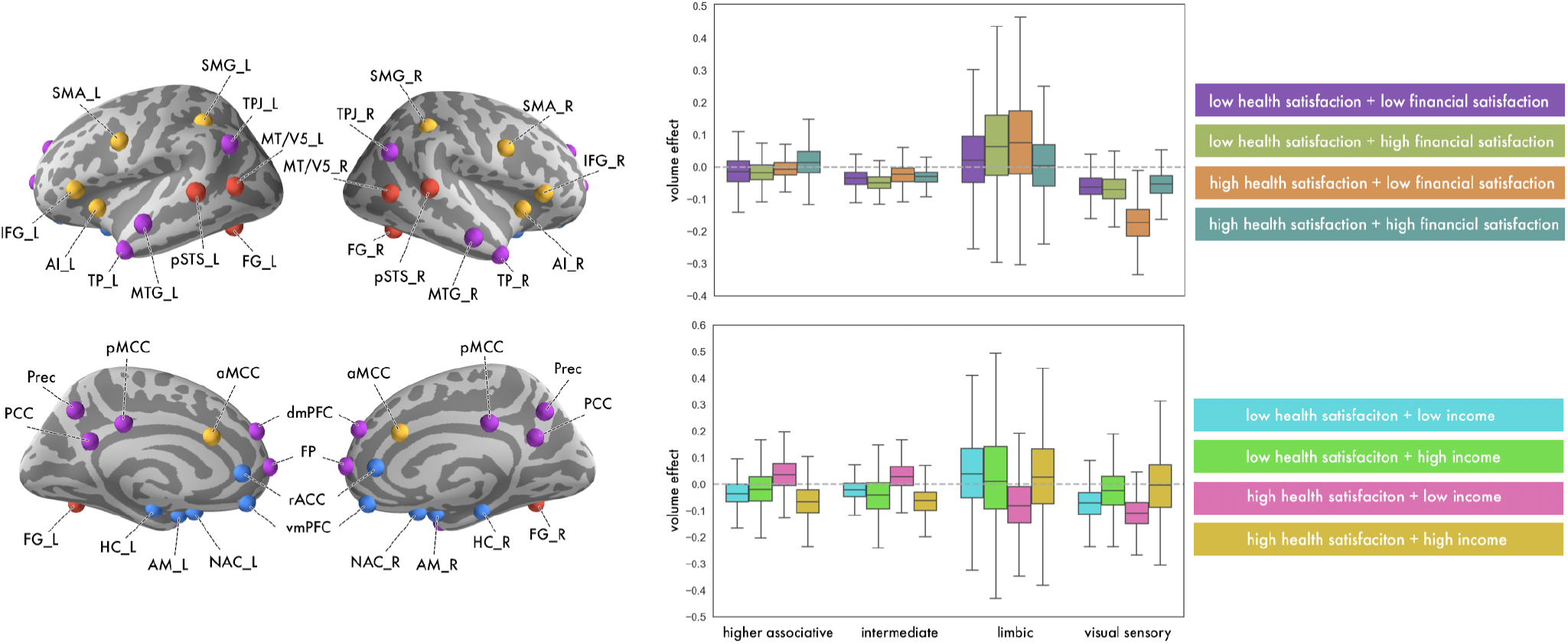
Network-wide health-related effects highlight the intermediate and visual-sensory systems. In addition to the within-network volume effects (Figs. 1–4), our hierarchical analysis framework explicitly modeled and identified network-wide volume effects as indicated by the inferred marginal posterior parameter distributions for the visual-sensory (red), intermediate (yellow), limbic (blue) and higher associative network (purple). Together, network-wide structural variation was most apparent in the intermediate network among participants content with their health, but of differing SES levels. In contrast, the visual-sensory network revealed income-related anatomical divergences among participants satisfied with their health. For abbreviations see Supplementary Table 1.

## Discussion

We investigated social brain variation in ~10,000 UK Biobank participants in an effort to understand how SES and health disparities may be reflected in grey matter volume. We tailored a Bayesian hierarchical modelling framework to capture coherent population-level correlates of health status and socioeconomic position. The two models separately analyzed the relationship between health satisfaction and SES in social brain structure: one model focused on income as an objective measure and the other model focused on financial satisfaction as a subjective measure. The identified brain-behavior associations showed region-specific and network-wide volume effects for all networks of the social brain atlas (Alcalá-López et al. 2018).

Interindividual differences in SES have previously been reported to influence health outcomes (Adler et al. 1994), and have also been linked to brain architecture (Gianaros et al. 2017). Our results thus confirm and extend these previous findings by providing neurobiological evidence in 10,000 individuals that discrepancies in health and SES markers can show diverging manifestations in social brain volume. As such, we observed volume effects both when measuring subjective and objective SES to show differentiation in grey matter volume in regions of the higher-associative network among individuals with a high health rating. Conversely, participants with a lower perceived quality of health showed distinct volume effects in regions of the limbic network as a function of household income. However, participants in good perceived health show selective income- and financial satisfaction-related deviations in grey matter volume mostly in visual-sensory network regions.

Our findings notably demonstrate the pervasiveness of SES-health disparities in social brain structure both at the single region and network level, as observed in both examined SES measures. Part of the higher associative network of the social brain, the dmPFC is known for its role in Theory-of-Mind (ToM) processes (e.g., Mitchell 2009). Such perspective-taking capacities are essential for moderating social interaction behavior by understanding the internal motivation, emotions, thoughts and beliefs of others (Carruthers and Smith 1996). A previous fMRI study investigated the neural mechanisms of social status, a construct that requires understanding the minds of others. The authors found that low-SES individuals may spontaneously consider the thoughts and feelings of others more often than high status individuals, as observed in higher dmPFC activation (Muscatell et al. 2012). The authors suggest the noted SES-based differences in dmPFC activation may be related to discrepancies in social perception. Indeed, a behavioral study found low-SES individuals to exhibit better empathic accuracy than individuals higher up on the social ladder (Kraus et al. 2010). As such, low-SES individuals may possess better skills than high-SES individuals at inferring the mental and emotional states of others (Muscatell et al. 2012).

Detailing these previous findings, we detected evidence for anatomical divergence in the dmPFC for individuals feeling unhealthy and unsatisfied with their financial situation compared to participants also feeling unhealthy, but happier with their finances. This deviation in dmPFC volume was also observed in the context of objective SES. Specifically, low-income participants showed the larger dmPFC volume effect, in both contexts of good and poor health status.

Furthermore, dmPFC involvement in complex social interaction behavior has been shown to have an impact on threat-related physiological responses (Eisenberger & Cole, 2012), which may bear relation with the dmPFC findings in our analyses. As previously evaluated by Eisenberger and colleagues, dmPFC activity may relate to alterations in cortisol activity from certain social stressors such as social exclusion (Eisenberger et al., 2007, 2011). In turn, these social stressors may be experienced as a threat to survival, and may affect downstream health by processes such as increasing the inflammatory response (Eisenberger & Cole, 2012). Together, our dmPFC findings in both indicators of socioeconomic lifestyle complement these studies that suggest lower SES individuals may read the minds of others better than higher-SES individuals, and may show sensitivity to negative social events.

Thinking about other individuals in the social environment also involves the TPJ, another brain region of our higher-associative network. Similar to the dmPFC, the TPJ is also linked to neurocognitive processes related to perspective-taking. In a previous fMRI study, the TPJ was repeatedly linked to perspective-taking during a series of false-belief stories tasks (Saxe and Kanwisher 2003). In our population-level results, dispersions in volume distributions were observed in the bilateral TPJ, especially for individuals in poor subjective health and dissatisfied with their finances. Lower-SES individuals have been noted to be more accurate at judging the emotions of others (Kraus et al. 2010). Kraus and colleagues speculate that low-SES individuals may focus more on external social situations to navigate and understand their life circumstances, compared to high-SES individuals. Hence, low-SES individuals may gravitate to others in their social surroundings for information, thus showing a SES-related pattern in perceiving and interacting with the social environment (Kraus et al. 2010).

In line with this contention, a structural MRI study assessed the relationship between exposure to socioeconomic disadvantage and brain structure (Gianaros et al. 2017). The authors found low SES, a composite measure of household poverty level, public assistance, education level, employment status and household income to be linked to reductions in grey matter volume in frontal and temporal regions extending into the TPJ and dmPFC (Gianaros et al. 2017). These reported reductions in cortical grey matter volume were also associated with poor health (Gianaros et al. 2017). In our population-level results, we observed participants with a lower perceived quality of health and unhappy with their financial situation to show the biggest volume effects in the TPJ, compared to the other subgroups in this model. Taken together, the identifid population volume deviations in the dmPFC and TPJ extend these previous findings by showing that neural manifestations of SES-health disparities are evident in regions of the social brain, which are commonly associated with perspective-taking.

Navigating the social environment also requires implicitly conforming to or explicitly knowing one’s place in society. A previous neuroimaging study investigated the neural processing of social hierarchies (Zink et al. 2008). The authors found activation in the somatosensory parts of the AI, a brain region belonging to our immediate network. In particular, AI activation was observed when high-status participants performed worse than an inferior player in an interactive game involving monetary reward (Zink et al. 2008). Neural activity changes in the AI also correlated with positive feelings associated with being of high social rank (Zink et al. 2008).

In human and non-human primates, social rank has been speculated to be deeply connected to long-term health outcomes, including consequences from stress (Sapolsky 2005). The connection between social rank and health has notably been observed in societies where instability exists in the social ordering of individuals, as reported from social hierarchies in monkeys (Sapolsky 2005). Our analyses in humans highlighted the AI to show select volume variation not only among participants with a lower health rating and low financial satisfaction, but also for participants of the same low health status, but earning a higher income. These variations in parameter volume distributions offer support for a link between low health status and AI volume, as a function of both objective and subjective SES. Indeed, Zink and colleagues attribute their findings in the AI to suggest a neural system that is pivotal for health-related risks associated with one’s anchoring in the hierarchical layers of society (Zink et al. 2008). Taken together, our results invite the speculation that the AI may be a neural correlate for social hierarchy processes, and may have volume adaptations related to social situations that can potentially threaten health outcomes.

Experiences in the social environment may not be the same at different positions in the social ladder of society. For example, factors that may influence the health status for low-SES individuals may not have the same impact for high-SES individuals, such as living conditions or adequate nutrition (Adler et al. 1994). Part of the intermediate network, SMG structure was found to be altered after nine months of cognitive mental training on tasks related to mindfulness, perspective-taking, and emotion (Valk et al., 2017). These structural plasticity findings in the SMG suggest sensitivity to changes in one’s social experience. In our study, we observed select volume effects in the SMG for participants with the same high health status, but in different socioeconomic milieus. Specifically, participants with high subjective health, but low financial satisfaction showed the larger volume effect compared to participants happier with their finances. However, high-income participants with high health satisfaction showed the larger volume effect compared to low-income participants also feeling healthy.

Despite participants having the same high health status, the observed SMG volume effects involving different SES levels may be related to one’s subjective experience in the social environment (Muscatell 2018). For example, recent evidence revealed a SES-based difference in social distancing behavior during the COVID-19 pandemic (Weill et al. 2020). In this mobile-data tracking study, social distancing varied as a function of neighborhood income level, with high-income neighborhoods being the most mobile before the pandemic compared to low-income neighborhoods. However, this trend reversed after the start of the pandemic, with low-income neighborhoods practicing less social distancing compared to high-income neighborhoods (Weill et al. 2020). The authors note that their findings provide support for the unequal distribution of SES-based disparities in health. Together with our present population results, these considerations hint at SMG involvement in SES-based subjective experiences in life circumstances.

Subjective social experiences such as feelings of subordination have also been shown to deteriorate one’s health (Sapolsky 2004). In our population results, the emotion-related AM showed strong objective and subjective SES-related volume effects, notably among the subgroups of participants expressing dissatisfaction with their health. At the heart of the limbic network of the social brain (Alcalá-López et al. 2018), the AM has previously been involved in responding to social threats, such as when viewing unpleasant facial expressions (Inagaki et al. 2012). In a previous fMRI study, participants were injected with a low dose of endotoxin, which induced a temporary physiological inflammatory response (Inagaki et al. 2012). Participants administered the endotoxin were found to show greater activation in the AM to images perceived as socially threatening compared to socially non-threatening images. This finding was not observed among participants administered the placebo.

Moreover, Muscatell and colleagues also observed activation in the AM from low-SES individuals in response to threatening faces during a social threat task (Muscatell et al. 2012). The activation to social threat in both of these brain-imaging studies suggests AM involvement in perceiving negative social situations in the social environment, especially for individuals low in socioeconomic standing or poor in health. Detailing these previous neuroimaging findings, we observed distinct volume variation in the AM in the context of low income and low financial satisfaction for the subgroups of participants with a low health rating. Taken together, our population-level findings suggest that the AM has a differentiated relation to varying levels of social standing and health status.

In addition to AM involvement in stressful social experiences, the HC, another region of the limbic system, has also been found to respond to threat and stress (Butterworth et al. 2012). Butterworth and colleagues suggest that experiencing chronic, stressful situations such as poor health and socioeconomic disadvantages may affect grey matter volume. This proposed association between stress and grey matter volume may relate to our observed volume effects in the HC among the subgroups of participants with a lower perceived health status. Indeed, Butterworth and colleagues found both HC and AM grey matter size to be smaller for participants who were more likely to experience socioeconomic hardships compared to participants with a better financial situation (Butterworth et al. 2012). Furthermore, the authors found financial hardship to also relate to depression and poor cognitive functioning (Butterworth et al. 2012). In relation to our population-level findings, we observed large volume effects in the HC among the subgroups of participants with a poor health rating in both contexts of subjective and objective SES. Thus, our results showing greater volume variation that is reliably associated with participants in poor health and experiencing socioeconomic disadvantage may have volume adaptations in the limbic AM and HC.

Moreover, our analysis highlighted financial satisfaction-related volume effects in the limbic NAC region among participants reporting high health satisfaction. Particularly, healthy participants expressing dissatisfaction with their finances showed the larger NAC volume effect compared to participants happier with their finances. As a major component of the ventral striatum, the NAC has previously been associated with monetary reward (Kable and Glimcher 2007), as well as social rewards (Rademacher et al. 2014). The NAC volume effects observed among our healthy subgroups may be closely linked to social status evaluation. For instance, a previous fMRI study showed NAC activation for both monetary rewards and acknowledgement of having a good reputation, thereby raising the possibility that monetary and social rewards share similar neural processes (Izuma et al. 2008). The authors suggest healthy individuals are motivated to seek social approval from others, and that a positive reputation may be a conditioned reward related to monetary gains.

Indeed, a previous fMRI study investigating the neural correlates of social hierarchy found NAC activation when participants viewed a socially superior individual during an interactive game compared to when viewing a socially inferior individual (Zink et al. 2008). This process may have downstream relations to long-term health, since high-SES individuals have been found to enjoy more positive health outcomes than low-SES individuals (Sapolsky 2004). Our findings thus invite the speculation that among our healthy subgroups, those who are unsatisfied with their finances may view themselves as inferior in status, which may have volume adaptations in the reward-related limbic NAC.

The subjective feeling of subordination may involve neural processes in regions at the lowest tier of the processing hierarchy. In brain regions of the visual-sensory network most known for social cue processing, our population results presented volume effects especially for the subgroups of participants satisfied with their health, in both SES measures. Notably, among these participants in good health, large structural deviations were observed in the pSTS, MT/V5 and FG regions in the context of both subjective and objective SES. Indeed, learning information about another person’s social status may be achieved through nonverbal cues in the social environment, such as through shifts in eye gaze (Pelphrey et al., 2004). A previous fMRI study found STS, FG and MT/V5 activation in a virtual reality eye gaze paradigm, where individuals either made eye contact with an approaching stranger or the stranger averted his eye gaze as he approached (Pelphrey et al., 2004). Notably, neural activity in the STS was most responsive when participants made eye contact with the approaching stranger than when he averted his gaze. The authors attribute this finding to the role of these visual-sensory regions in processing social information from biological motion, and highlight the sensitivity of the STS to nonverbal social cues.

In our visual-sensory network region findings, we observed the strongest volume effects in the bilateral pSTS among participants feeling healthy, notably in the context of financial satisfaction. In line with this finding, a previous fMRI study found greater pSTS and mPFC activation when participants made judgments about another individual’s status compared to judgments about another person’s physical appearance (Mason et al. 2014). The authors suggest that these regions are implicated in mentalizing about other individuals, and even more so when assessing their social status (Mason et al. 2014). Furthermore, humans avert their gaze from others they perceive to be dominant, thus signaling submission (Holland et al. 2017). Together, our results offer support for these previous neuroimaging findings that humans in good health may be able to detect differences in social status in brain regions at the lowest tier of the processing hierarchy.

## Conclusion

The link between socioeconomic status and health disparity is deeply rooted in human societies. Previous research has coalesced in the conclusion that the “well off” typically experience fewer detrimental health outcomes compared to individuals with lower position on the social ladder (Adler et al. 1994). Using a fully probabilistic modelling strategy in a population cohort of 10,000 individuals, we uncovered patterns of similarity and divergence between health and two measures of SES in the social brain, especially in regions of the visual-sensory and higher associative networks. In this way, we provide a population-level perspective into the brain association of health disparity experienced at different subjective and objective socioeconomic levels. Our observations contribute to further disentangling the complex interplay of health disparities and socioeconomic standing at the neurobiological level.

## Acknowledgements

D.B. has been funded by the Brain Canada Foundation, through the Canada Brain Research Fund, with the financial support of Health Canada, National Institutes of Health (NIH R01 AG068563A), the Canadian Institute of Health Research (CIHR 438531), the Healthy Brains Healthy Lives initiative (Canada First Research Excellence fund), Google (Research Award, Teaching Award), and by the CIFAR Artificial Intelligence Chairs program (Canada Institute for Advanced Research).

## Supplementary Material

**Supplementary Table 1.**
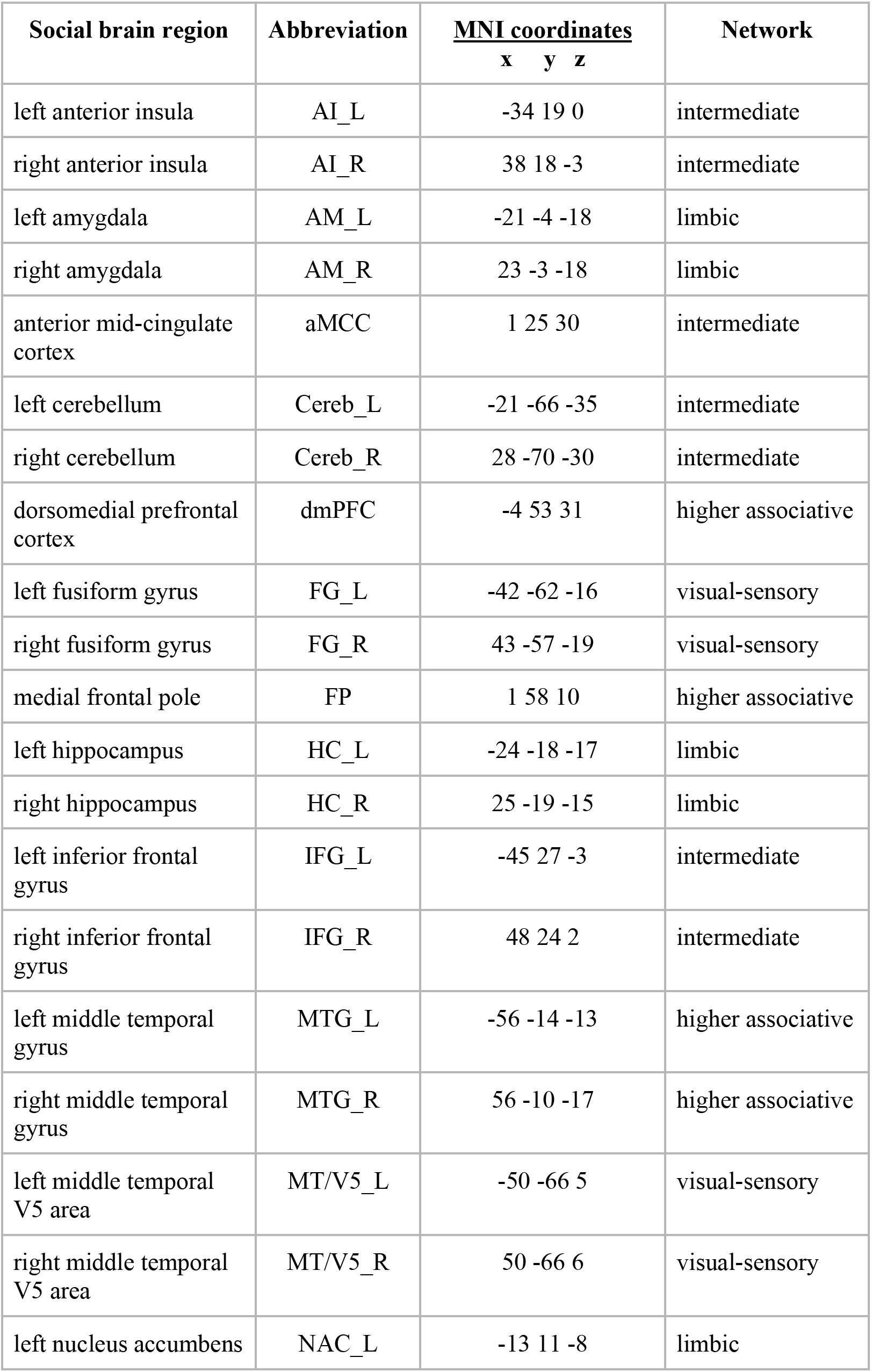
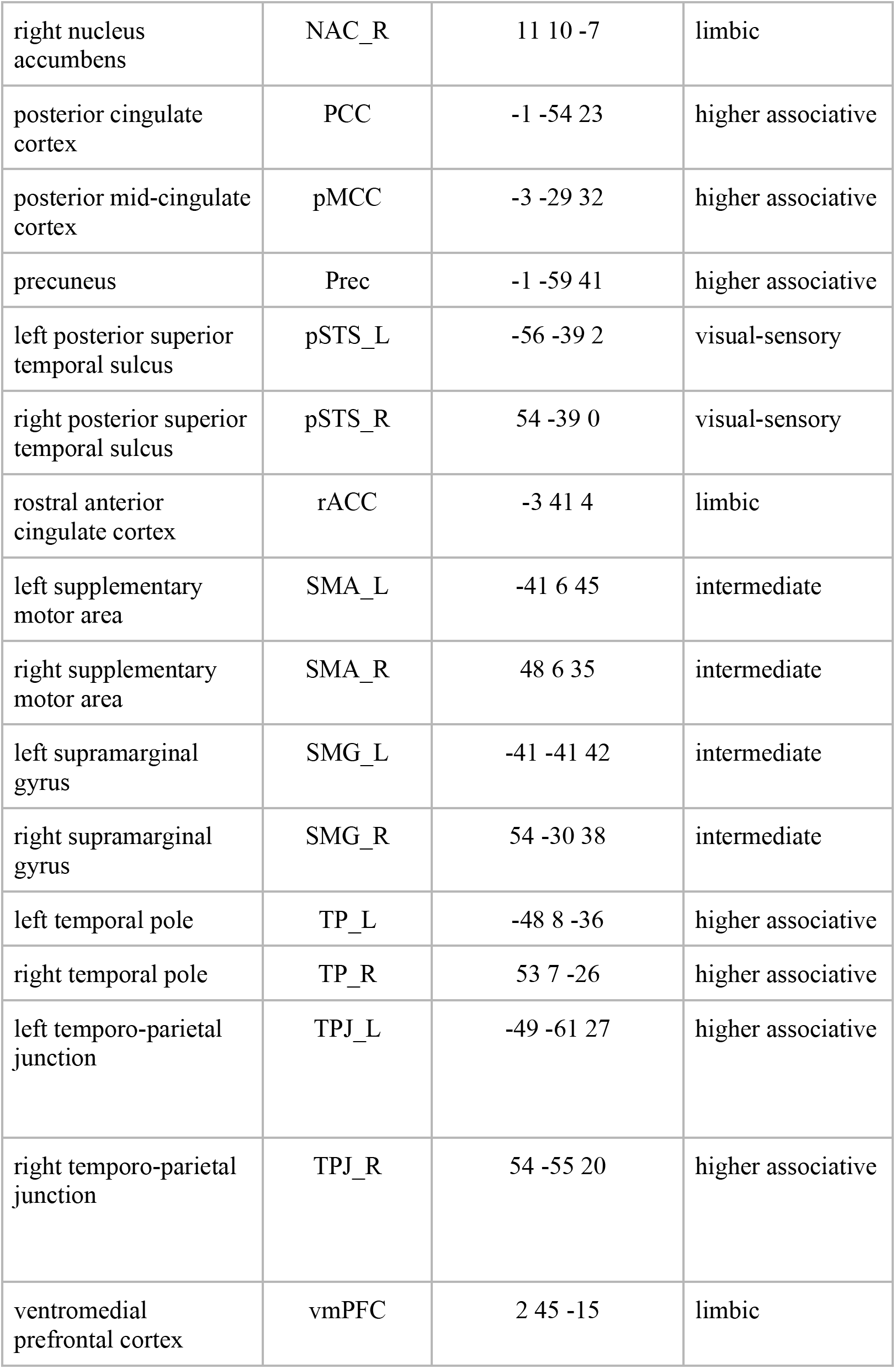
Social brain atlas regions and their MNI coordinates. Social brain regions and their respective functional network as depicted in (Alcalá-López et al., 2018).

| **Social brain region** | **Abbreviation** | **<u>MNI coordinates</u>**<br>**x y z** | **Network** |
| --- | --- | --- | --- |
| left anterior insula | AI_L | -34 19 0 | intermediate |
| right anterior insula | AI_R | 38 18 -3 | intermediate |
| left amygdala | AM_L | -21 -4 -18 | limbic |
| right amygdala | AM_R | 23 -3 -18 | limbic |
| anterior mid-cingulate cortex | aMCC | 1 25 30 | intermediate |
| left cerebellum | Cereb_L | -21 -66 -35 | intermediate |
| right cerebellum | Cereb_R | 28 -70 -30 | intermediate |
| dorsomedial prefrontal cortex | dmPFC | -4 53 31 | higher associative |
| left fusiform gyrus | FG_L | -42 -62 -16 | visual-sensory |
| right fusiform gyrus | FG_R | 43 -57 -19 | visual-sensory |
| medial frontal pole | FP | 1 58 10 | higher associative |
| left hippocampus | HC_L | -24 -18 -17 | limbic |
| right hippocampus | HC_R | 25 -19 -15 | limbic |
| left inferior frontal gyrus | IFG_L | -45 27 -3 | intermediate |
| right inferior frontal gyrus | IFG_R | 48 24 2 | intermediate |
| left middle temporal gyrus | MTG_L | -56 -14 -13 | higher associative |
| right middle temporal gyrus | MTG_R | 56 -10 -17 | higher associative |
| left middle temporal V5 area | MT/V5_L | -50 -66 5 | visual-sensory |
| right middle temporal V5 area | MT/V5_R | 50 -66 6 | visual-sensory |
| left nucleus accumbens | NAC_L | -13 11 -8 | limbic |
| right nucleus accumbens | NAC_R | 11 10 -7 | limbic |
| posterior cingulate cortex | PCC | -1 -54 23 | higher associative |
| posterior mid-cingulate cortex | pMCC | -3 -29 32 | higher associative |
| precuneus | Prec | -1 -59 41 | higher associative |
| left posterior superior temporal sulcus | pSTS_L | -56 -39 2 | visual-sensory |
| right posterior superior temporal sulcus | pSTS_R | 54 -39 0 | visual-sensory |
| rostral anterior cingulate cortex | rACC | -3 41 4 | limbic |
| left supplementary motor area | SMA_L | -41 6 45 | intermediate |
| right supplementary motor area | SMA_R | 48 6 35 | intermediate |
| left supramarginal gyrus | SMG_L | -41 -41 42 | intermediate |
| right supramarginal gyrus | SMG_R | 54 -30 38 | intermediate |
| left temporal pole | TP_L | -48 8 -36 | higher associative |
| right temporal pole | TP_R | 53 7 -26 | higher associative |
| left temporo-parietal junction | TPJ_L | -49 -61 27 | higher associative |
| right temporo-parietal junction | TPJ_R | 54 -55 20 | higher associative |
| ventromedial prefrontal cortex | vmPFC | 2 45 -15 | limbic |

**Supplementary Table 2.** HPDI and posterior mean parameter values for health satisfaction and financial satisfaction.

| Health satisfaction + financial satisfaction |  |  |  |  |
| --- | --- | --- | --- | --- |
| subgroup | higher_HPDI | lower_HPDI | posterior_mean | roi |
| Unhappy health + unhappy financially | 0.163 | -0.064 | 0.048 | FG_L |
| Unhappy health + happy financially | 0.147 | -0.106 | 0.005 | FG_L |
| Happy health + unhappy financially | 0.191 | -0.163 | 0.000 | FG_L |
| Happy health + happy financially | 0.108 | -0.106 | -0.008 | FG_L |
| Unhappy health + unhappy financially | -0.019 | -0.223 | -0.110 | FG_R |
| Unhappy health + happy financially | -0.021 | -0.272 | -0.130 | FG_R |
| Happy health + unhappy financially | -0.092 | -0.364 | -0.220 | FG_R |
| Happy health + happy financially | 0.079 | -0.128 | -0.032 | FG_R |
| Unhappy health + unhappy financially | 0.012 | -0.157 | -0.069 | pSTS_L |
| Unhappy health + happy financially | -0.005 | -0.226 | -0.109 | pSTS_L |
| Happy health + unhappy financially | -0.122 | -0.408 | -0.259 | pSTS_L |
| Happy health + happy financially | -0.012 | -0.214 | -0.105 | pSTS_L |
| Unhappy health + unhappy financially | 0.021 | -0.167 | -0.072 | pSTS_R |
| Unhappy health + happy financially | -0.019 | -0.237 | -0.115 | pSTS_R |
| Happy health + unhappy financially | -0.122 | -0.375 | -0.242 | pSTS_R |
| Happy health + happy financially | 0.034 | -0.180 | -0.074 | pSTS_R |
| Unhappy health + unhappy financially | 0.032 | -0.141 | -0.058 | MTV5_L |
| Unhappy health + happy financially | 0.072 | -0.137 | -0.040 | MTV5_L |
| Happy health + unhappy financially | -0.040 | -0.250 | -0.147 | MTV5_L |
| Happy health + happy financially | 0.122 | -0.064 | 0.024 | MTV5_L |
| Unhappy health + unhappy financially | -0.024 | -0.185 | -0.101 | MTV5_R |
| Unhappy health + happy financially | 0.090 | -0.112 | -0.018 | MTV5_R |
| Happy health + unhappy financially | -0.062 | -0.283 | -0.170 | MTV5_R |
| Happy health + happy financially | -0.042 | -0.231 | -0.132 | MTV5_R |
| Unhappy health + unhappy financially | 0.346 | 0.036 | 0.189 | AM_L |
| Unhappy health + happy financially | 0.345 | -0.075 | 0.132 | AM_L |
| Happy health + unhappy financially | 0.112 | -0.331 | -0.112 | AM_L |
| Happy health + happy financially | 0.147 | -0.192 | -0.016 | AM_L |
| Unhappy health + unhappy financially | 0.620 | 0.173 | 0.387 | AM_R |
| Unhappy health + happy financially | 1.073 | 0.377 | 0.721 | AM_R |
| Happy health + unhappy financially | 1.412 | 0.676 | 1.036 | AM_R |
| Happy health + happy financially | 0.578 | 0.067 | 0.306 | AM_R |
| Unhappy health + unhappy financially | 0.468 | 0.043 | 0.258 | HC_L |
| Unhappy health + happy financially | 0.661 | 0.136 | 0.397 | HC_L |
| Happy health + unhappy financially | 0.567 | -0.019 | 0.269 | HC_L |
| Happy health + happy financially | 0.413 | -0.033 | 0.188 | HC_L |
| Unhappy health + unhappy financially | -0.085 | -0.436 | -0.256 | HC_R |
| Unhappy health + happy financially | 0.008 | -0.458 | -0.229 | HC_R |
| Happy health + unhappy financially | 0.203 | -0.297 | -0.047 | HC_R |
| Happy health + happy financially | 0.256 | -0.152 | 0.061 | HC_R |
| Unhappy health + unhappy financially | 0.119 | -0.099 | 0.006 | vmPFC |
| Unhappy health + happy financially | 0.082 | -0.201 | -0.060 | vmPFC |
| Happy health + unhappy financially | 0.136 | -0.199 | -0.028 | vmPFC |
| Happy health + happy financially | 0.046 | -0.189 | -0.072 | vmPFC |
| Unhappy health + unhappy financially | -0.276 | -0.495 | -0.389 | NAC_L |
| Unhappy health + happy financially | -0.260 | -0.558 | -0.413 | NAC_L |
| Happy health + unhappy financially | -0.015 | -0.328 | -0.172 | NAC_L |
| Happy health + happy financially | -0.255 | -0.495 | -0.376 | NAC_L |
| Unhappy health + unhappy financially | 0.151 | -0.114 | 0.028 | NAC_R |
| Unhappy health + happy financially | 0.232 | -0.132 | 0.048 | NAC_R |
| Happy health + unhappy financially | -0.073 | -0.429 | -0.252 | NAC_R |
| Happy health + happy financially | 0.160 | -0.130 | 0.013 | NAC_R |
| Unhappy health + unhappy financially | 0.092 | -0.115 | -0.014 | rACC |
| Unhappy health + happy financially | 0.086 | -0.169 | -0.040 | rACC |
| Happy health + unhappy financially | 0.028 | -0.240 | -0.103 | rACC |
| Happy health + happy financially | 0.035 | -0.175 | -0.067 | rACC |
| Unhappy health + unhappy financially | -0.029 | -0.243 | -0.128 | aMCC |
| Unhappy health + happy financially | -0.031 | -0.268 | -0.134 | aMCC |
| Happy health + unhappy financially | 0.048 | -0.181 | -0.065 | aMCC |
| Happy health + happy financially | 0.024 | -0.141 | -0.051 | aMCC |
| Unhappy health + unhappy financially | 0.026 | -0.184 | -0.079 | AI_L |
| Unhappy health + happy financially | 0.023 | -0.189 | -0.075 | AI_L |
| Happy health + unhappy financially | 0.102 | -0.142 | -0.024 | AI_L |
| Happy health + happy financially | 0.096 | -0.105 | -0.014 | AI_L |
| Unhappy health + unhappy financially | 0.279 | -0.012 | 0.122 | AI_R |
| Unhappy health + happy financially | 0.143 | -0.108 | -0.006 | AI_R |
| Happy health + unhappy financially | 0.171 | -0.101 | 0.026 | AI_R |
| Happy health + happy financially | 0.182 | -0.064 | 0.036 | AI_R |
| Unhappy health + unhappy financially | 0.074 | -0.065 | 0.002 | SMG_L |
| Unhappy health + happy financially | 0.043 | -0.116 | -0.039 | SMG_L |
| Happy health + unhappy financially | 0.089 | -0.095 | -0.003 | SMG_L |
| Happy health + happy financially | 0.056 | -0.092 | -0.023 | SMG_L |
| Unhappy health + unhappy financially | 0.057 | -0.087 | -0.015 | SMG_R |
| Unhappy health + happy financially | 0.105 | -0.071 | 0.009 | SMG_R |
| Happy health + unhappy financially | 0.176 | -0.032 | 0.065 | SMG_R |
| Happy health + happy financially | 0.076 | -0.081 | -0.007 | SMG_R |
| Unhappy health + unhappy financially | 0.033 | -0.130 | -0.049 | SMA_L |
| Unhappy health + happy financially | 0.068 | -0.137 | -0.043 | SMA_L |
| Happy health + unhappy financially | 0.018 | -0.195 | -0.085 | SMA_L |
| Happy health + happy financially | -0.008 | -0.182 | -0.088 | SMA_L |
| Unhappy health + unhappy financially | 0.075 | -0.104 | -0.018 | SMA_R |
| Unhappy health + happy financially | 0.011 | -0.180 | -0.079 | SMA_R |
| Happy health + unhappy financially | 0.041 | -0.165 | -0.057 | SMA_R |
| Happy health + happy financially | 0.021 | -0.145 | -0.057 | SMA_R |
| Unhappy health + unhappy financially | 0.007 | -0.166 | -0.080 | IFG_L |
| Unhappy health + happy financially | 0.044 | -0.147 | -0.052 | IFG_L |
| Happy health + unhappy financially | 0.007 | -0.218 | -0.098 | IFG_L |
| Happy health + happy financially | -0.008 | -0.193 | -0.090 | IFG_L |
| Unhappy health + unhappy financially | -0.013 | -0.185 | -0.095 | IFG_R |
| Unhappy health + happy financially | 0.016 | -0.164 | -0.070 | IFG_R |
| Happy health + unhappy financially | 0.031 | -0.206 | -0.074 | IFG_R |
| Happy health + happy financially | 0.039 | -0.123 | -0.039 | IFG_R |
| Unhappy health + unhappy financially | 0.084 | -0.129 | -0.022 | Cereb_L |
| Unhappy health + happy financially | 0.073 | -0.149 | -0.036 | Cereb_L |
| Happy health + unhappy financially | 0.268 | -0.033 | 0.105 | Cereb_L |
| Happy health + happy financially | 0.142 | -0.075 | 0.012 | Cereb_L |
| Unhappy health + unhappy financially | 0.122 | -0.091 | 0.009 | Cereb_R |
| Unhappy health + happy financially | 0.118 | -0.106 | -0.004 | Cereb_R |
| Happy health + unhappy financially | 0.076 | -0.196 | -0.043 | Cereb_R |
| Happy health + happy financially | 0.070 | -0.119 | -0.019 | Cereb_R |
| Unhappy health + unhappy financially | 0.024 | -0.122 | -0.046 | FP |
| Unhappy health + happy financially | -0.002 | -0.187 | -0.090 | FP |
| Happy health + unhappy financially | 0.034 | -0.146 | -0.050 | FP |
| Happy health + happy financially | -0.002 | -0.178 | -0.092 | FP |
| Unhappy health + unhappy financially | 0.008 | -0.144 | -0.067 | dmPFC |
| Unhappy health + happy financially | 0.131 | -0.063 | 0.030 | dmPFC |
| Happy health + unhappy financially | 0.091 | -0.096 | -0.004 | dmPFC |
| Happy health + happy financially | 0.150 | -0.033 | 0.058 | dmPFC |
| Unhappy health + unhappy financially | 0.123 | -0.134 | -0.001 | PCC |
| Unhappy health + happy financially | 0.056 | -0.243 | -0.085 | PCC |
| Happy health + unhappy financially | 0.252 | -0.047 | 0.069 | PCC |
| Happy health + happy financially | 0.052 | -0.233 | -0.081 | PCC |
| Unhappy health + unhappy financially | -0.089 | -0.258 | -0.173 | TPJ_L |
| Unhappy health + happy financially | 0.015 | -0.179 | -0.081 | TPJ_L |
| Happy health + unhappy financially | 0.044 | -0.155 | -0.050 | TPJ_L |
| Happy health + happy financially | -0.070 | -0.260 | -0.166 | TPJ_L |
| Unhappy health + unhappy financially | -0.045 | -0.221 | -0.135 | TPJ_R |
| Unhappy health + happy financially | 0.069 | -0.148 | -0.041 | TPJ_R |
| Happy health + unhappy financially | 0.057 | -0.129 | -0.033 | TPJ_R |
| Happy health + happy financially | 0.091 | -0.114 | -0.012 | TPJ_R |
| Unhappy health + unhappy financially | 0.029 | -0.151 | -0.057 | Prec |
| Unhappy health + happy financially | 0.028 | -0.179 | -0.075 | Prec |
| Happy health + unhappy financially | 0.079 | -0.122 | -0.018 | Prec |
| Happy health + happy financially | 0.154 | -0.045 | 0.056 | Prec |
| Unhappy health + unhappy financially | -0.004 | -0.175 | -0.085 | MTG_L |
| Unhappy health + happy financially | 0.076 | -0.131 | -0.023 | MTG_L |
| Happy health + unhappy financially | 0.049 | -0.154 | -0.040 | MTG_L |
| Happy health + happy financially | 0.055 | -0.161 | -0.053 | MTG_L |
| Unhappy health + unhappy financially | -0.021 | -0.220 | -0.119 | MTG_R |
| Unhappy health + happy financially | 0.032 | -0.204 | -0.078 | MTG_R |
| Happy health + unhappy financially | 0.088 | -0.105 | -0.011 | MTG_R |
| Happy health + happy financially | 0.033 | -0.199 | -0.080 | MTG_R |
| Unhappy health + unhappy financially | 0.302 | 0.057 | 0.176 | TP_L |
| Unhappy health + happy financially | 0.211 | -0.061 | 0.064 | TP_L |
| Happy health + unhappy financially | 0.183 | -0.071 | 0.040 | TP_L |
| Happy health + happy financially | 0.288 | 0.036 | 0.159 | TP_L |
| Unhappy health + unhappy financially | 0.207 | 0.003 | 0.105 | TP_R |
| Unhappy health + happy financially | 0.150 | -0.078 | 0.034 | TP_R |
| Happy health + unhappy financially | 0.072 | -0.146 | -0.033 | TP_R |
| Happy health + happy financially | 0.159 | -0.065 | 0.043 | TP_R |
| Unhappy health + unhappy financially | 0.362 | 0.102 | 0.230 | pMCC |
| Unhappy health + happy financially | 0.350 | -0.010 | 0.153 | pMCC |
| Happy health + unhappy financially | 0.245 | -0.048 | 0.066 | pMCC |
| Happy health + happy financially | 0.487 | 0.168 | 0.322 | pMCC |

**Supplementary Table 3.** HPDI and posterior mean parameter values for income and health satisfaction.

| Health satisfaction + income |  |  |  |  |
| --- | --- | --- | --- | --- |
| group | higher_HPDI | lower_HPDI | posterior_mean | roi |
| Unhappy health + low income | 0.232 | -0.164 | -0.001 | FG_L |
| Unhappy health + high income | 0.339 | -0.220 | 0.027 | FG_L |
| Happy health + low income | 0.114 | -0.244 | -0.079 | FG_L |
| Happy health + high income | 0.305 | -0.372 | -0.034 | FG_L |
| Unhappy health + low income | 0.218 | -0.162 | 0.004 | FG_R |
| Unhappy health + high income | 0.249 | -0.345 | -0.029 | FG_R |
| Happy health + low income | 0.112 | -0.253 | -0.086 | FG_R |
| Happy health + high income | 0.390 | -0.377 | 0.006 | FG_R |
| Unhappy health + low income | 0.019 | -0.309 | -0.126 | pSTS_L |
| Unhappy health + high income | 0.251 | -0.323 | -0.024 | pSTS_L |
| Happy health + low income | 0.090 | -0.245 | -0.092 | pSTS_L |
| Happy health + high income | 0.146 | -0.672 | -0.212 | pSTS_L |
| Unhappy health + low income | 0.062 | -0.252 | -0.087 | pSTS_R |
| Unhappy health + high income | 0.249 | -0.262 | -0.016 | pSTS_R |
| Happy health + low income | 0.006 | -0.372 | -0.163 | pSTS_R |
| Happy health + high income | 0.485 | -0.311 | 0.058 | pSTS_R |
| Unhappy health + low income | 0.087 | -0.202 | -0.063 | MTV5_L |
| Unhappy health + high income | 0.278 | -0.196 | 0.014 | MTV5_L |
| Happy health + low income | 0.061 | -0.251 | -0.090 | MTV5_L |
| Happy health + high income | 0.432 | -0.117 | 0.152 | MTV5_L |
| Unhappy health + low income | -0.020 | -0.308 | -0.157 | MTV5_R |
| Unhappy health + high income | 0.115 | -0.402 | -0.106 | MTV5_R |
| Happy health + low income | -0.013 | -0.328 | -0.155 | MTV5_R |
| Happy health + high income | 0.315 | -0.283 | 0.018 | MTV5_R |
| Unhappy health + low income | 0.268 | -0.430 | -0.071 | AM_L |
| Unhappy health + high income | 0.665 | -0.186 | 0.186 | AM_L |
| Happy health + low income | 0.148 | -0.413 | -0.124 | AM_L |
| Happy health + high income | 0.203 | -0.882 | -0.270 | AM_L |
| Unhappy health + low income | 1.329 | 0.212 | 0.751 | AM_R |
| Unhappy health + high income | 1.057 | -0.325 | 0.311 | AM_R |
| Happy health + low income | 0.636 | -0.196 | 0.173 | AM_R |
| Happy health + high income | 0.862 | -0.368 | 0.202 | AM_R |
| Unhappy health + low income | 0.260 | -0.478 | -0.110 | HC_L |
| Unhappy health + high income | 1.109 | -0.250 | 0.326 | HC_L |
| Happy health + low income | 0.515 | -0.247 | 0.104 | HC_L |
| Happy health + high income | 1.525 | -0.046 | 0.642 | HC_L |
| Unhappy health + low income | 0.376 | -0.278 | 0.054 | HC_R |
| Unhappy health + high income | 0.207 | -1.290 | -0.408 | HC_R |
| Happy health + low income | 0.287 | -0.386 | -0.042 | HC_R |
| Happy health + high income | 0.658 | -0.581 | 0.057 | HC_R |
| Unhappy health + low income | 0.186 | -0.265 | -0.041 | vmPFC |
| Unhappy health + high income | 0.448 | -0.394 | 0.059 | vmPFC |
| Happy health + low income | 0.148 | -0.367 | -0.109 | vmPFC |
| Happy health + high income | 0.401 | -0.463 | -0.037 | vmPFC |
| Unhappy health + low income | -0.133 | -0.562 | -0.348 | NAC_L |
| Unhappy health + high income | -0.056 | -1.410 | -0.605 | NAC_L |
| Happy health + low income | -0.107 | -0.634 | -0.364 | NAC_L |
| Happy health + high income | 0.232 | -0.559 | -0.157 | NAC_L |
| Unhappy health + low income | 0.324 | -0.142 | 0.091 | NAC_R |
| Unhappy health + high income | 0.779 | -0.365 | 0.136 | NAC_R |
| Happy health + low income | 0.103 | -0.458 | -0.182 | NAC_R |
| Happy health + high income | 0.314 | -0.622 | -0.135 | NAC_R |
| Unhappy health + low income | 0.187 | -0.177 | 0.010 | rACC |
| Unhappy health + high income | 0.572 | -0.109 | 0.214 | rACC |
| Happy health + low income | 0.151 | -0.290 | -0.072 | rACC |
| Happy health + high income | 0.327 | -0.289 | 0.012 | rACC |
| Unhappy health + low income | 0.135 | -0.137 | -0.007 | aMCC |
| Unhappy health + high income | 0.154 | -0.564 | -0.161 | aMCC |
| Happy health + low income | 0.221 | -0.149 | 0.033 | aMCC |
| Happy health + high income | 0.155 | -0.273 | -0.064 | aMCC |
| Unhappy health + low income | 0.157 | -0.161 | -0.010 | AI_L |
| Unhappy health + high income | 0.070 | -0.701 | -0.255 | AI_L |
| Happy health + low income | 0.224 | -0.215 | 0.018 | AI_L |
| Happy health + high income | 0.219 | -0.254 | -0.051 | AI_L |
| Unhappy health + low income | 0.188 | -0.141 | 0.006 | AI_R |
| Unhappy health + high income | 0.432 | -0.318 | 0.034 | AI_R |
| Happy health + low income | 0.459 | -0.025 | 0.182 | AI_R |
| Happy health + high income | 0.139 | -0.349 | -0.084 | AI_R |
| Unhappy health + low income | 0.208 | -0.076 | 0.046 | SMG_L |
| Unhappy health + high income | 0.430 | -0.096 | 0.147 | SMG_L |
| Happy health + low income | 0.190 | -0.127 | 0.033 | SMG_L |
| Happy health + high income | 0.206 | -0.184 | -0.017 | SMG_L |
| Unhappy health + low income | 0.158 | -0.097 | 0.023 | SMG_R |
| Unhappy health + high income | 0.341 | -0.130 | 0.094 | SMG_R |
| Happy health + low income | 0.275 | -0.065 | 0.096 | SMG_R |
| Happy health + high income | 0.054 | -0.320 | -0.105 | SMG_R |
| Unhappy health + low income | 0.113 | -0.148 | -0.015 | SMA_L |
| Unhappy health + high income | 0.278 | -0.280 | -0.008 | SMA_L |
| Happy health + low income | 0.081 | -0.349 | -0.109 | SMA_L |
| Happy health + high income | 0.148 | -0.240 | -0.059 | SMA_L |
| Unhappy health + low income | 0.022 | -0.315 | -0.120 | SMA_R |
| Unhappy health + high income | 0.135 | -0.493 | -0.151 | SMA_R |
| Happy health + low income | 0.133 | -0.202 | -0.019 | SMA_R |
| Happy health + high income | 0.128 | -0.268 | -0.065 | SMA_R |
| Unhappy health + low income | 0.069 | -0.203 | -0.055 | IFG_L |
| Unhappy health + high income | 0.223 | -0.298 | -0.032 | IFG_L |
| Happy health + low income | 0.114 | -0.274 | -0.062 | IFG_L |
| Happy health + high income | 0.104 | -0.307 | -0.082 | IFG_L |
| Unhappy health + low income | 0.051 | -0.202 | -0.064 | IFG_R |
| Unhappy health + high income | 0.278 | -0.283 | -0.014 | IFG_R |
| Happy health + low income | 0.156 | -0.228 | -0.028 | IFG_R |
| Happy health + high income | 0.108 | -0.346 | -0.087 | IFG_R |
| Unhappy health + low income | 0.142 | -0.147 | -0.010 | Cereb_L |
| Unhappy health + high income | 0.229 | -0.525 | -0.108 | Cereb_L |
| Happy health + low income | 0.391 | -0.024 | 0.151 | Cereb_L |
| Happy health + high income | 0.128 | -0.250 | -0.059 | Cereb_L |
| Unhappy health + low income | 0.105 | -0.170 | -0.032 | Cereb_R |
| Unhappy health + high income | 0.357 | -0.343 | -0.005 | Cereb_R |
| Happy health + low income | 0.241 | -0.133 | 0.055 | Cereb_R |
| Happy health + high income | 0.223 | -0.205 | -0.025 | Cereb_R |
| Unhappy health + low income | 0.068 | -0.209 | -0.071 | FP |
| Unhappy health + high income | 0.196 | -0.297 | -0.047 | FP |
| Happy health + low income | 0.047 | -0.363 | -0.143 | FP |
| Happy health + high income | 0.096 | -0.330 | -0.103 | FP |
| Unhappy health + low income | 0.025 | -0.273 | -0.120 | dmPFC |
| Unhappy health + high income | 0.260 | -0.243 | -0.003 | dmPFC |
| Happy health + low income | 0.294 | -0.079 | 0.097 | dmPFC |
| Happy health + high income | 0.238 | -0.222 | -0.013 | dmPFC |
| Unhappy health + low income | 0.218 | -0.206 | 0.003 | PCC |
| Unhappy health + high income | 0.349 | -0.286 | 0.009 | PCC |
| Happy health + low income | 0.316 | -0.170 | 0.072 | PCC |
| Happy health + high income | 0.251 | -0.317 | -0.056 | PCC |
| Unhappy health + low income | 0.032 | -0.278 | -0.118 | TPJ_L |
| Unhappy health + high income | 0.098 | -0.412 | -0.134 | TPJ_L |
| Happy health + low income | 0.061 | -0.343 | -0.133 | TPJ_L |
| Happy health + high income | 0.116 | -0.349 | -0.100 | TPJ_L |
| Unhappy health + low income | 0.036 | -0.270 | -0.114 | TPJ_R |
| Unhappy health + high income | 0.057 | -0.550 | -0.193 | TPJ_R |
| Happy health + low income | 0.194 | -0.176 | 0.010 | TPJ_R |
| Happy health + high income | 0.176 | -0.294 | -0.052 | TPJ_R |
| Unhappy health + low income | 0.147 | -0.164 | -0.011 | Prec |
| Unhappy health + high income | 0.141 | -0.449 | -0.107 | Prec |
| Happy health + low income | 0.282 | -0.128 | 0.074 | Prec |
| Happy health + high income | 0.216 | -0.231 | -0.029 | Prec |
| Unhappy health + low income | 0.075 | -0.238 | -0.073 | MTG_L |
| Unhappy health + high income | 0.353 | -0.246 | 0.032 | MTG_L |
| Happy health + low income | 0.221 | -0.178 | 0.030 | MTG_L |
| Happy health + high income | 0.213 | -0.240 | -0.033 | MTG_L |
| Unhappy health + low income | 0.088 | -0.264 | -0.079 | MTG_R |
| Unhappy health + high income | 0.262 | -0.338 | -0.032 | MTG_R |
| Happy health + low income | 0.273 | -0.186 | 0.050 | MTG_R |
| Happy health + high income | 0.061 | -0.464 | -0.153 | MTG_R |
| Unhappy health + low income | 0.326 | -0.091 | 0.095 | TP_L |
| Unhappy health + high income | 0.328 | -0.281 | 0.024 | TP_L |
| Happy health + low income | 0.325 | -0.122 | 0.097 | TP_L |
| Happy health + high income | 0.191 | -0.302 | -0.055 | TP_L |
| Unhappy health + low income | 0.116 | -0.223 | -0.050 | TP_R |
| Unhappy health + high income | 0.494 | -0.123 | 0.141 | TP_R |
| Happy health + low income | 0.212 | -0.216 | 0.006 | TP_R |
| Happy health + high income | 0.164 | -0.333 | -0.075 | TP_R |
| Unhappy health + low income | 0.484 | -0.061 | 0.197 | pMCC |
| Unhappy health + high income | 0.548 | -0.176 | 0.114 | pMCC |
| Happy health + low income | 0.566 | 0.014 | 0.256 | pMCC |
| Happy health + high income | 0.232 | -0.281 | -0.035 | pMCC |

